# RNAi screen reveals a role for PACSIN2 and caveolins during bacterial cell-to-cell spread

**DOI:** 10.1101/599795

**Authors:** Allen G. Sanderlin, Cassandra Vondrak, Arianna J. Scricco, Indro Fedrigo, Vida Ahyong, Rebecca L. Lamason

## Abstract

*Listeria monocytogenes* is a human bacterial pathogen that disseminates through host tissues using a process called cell-to-cell spread. This critical yet understudied virulence strategy resembles a vesicular form of intercellular trafficking that allows *L. monocytogenes* to move between host cells without escaping the cell. Interestingly, eukaryotic cells can also directly exchange cellular components via intercellular communication pathways (e.g. trans-endocytosis) using cell-cell adhesion, membrane trafficking, and membrane remodeling proteins. Therefore, we hypothesized that *L. monocytogenes* would hijack these types of host proteins during spread. Using a focused RNAi screen, we identified 22 host genes that are important for *L. monocytogenes* spread. We then found that caveolins (CAV1 and CAV2) and the membrane sculpting F-BAR protein PACSIN2 promote *L. monocytogenes* protrusion engulfment during spread, and that PACSIN2 specifically localized to protrusions. Overall, our study demonstrates that host intercellular communication pathways may be co-opted during bacterial spread and that specific trafficking and membrane remodeling proteins promote bacterial protrusion resolution.

**Summary:** The human bacterial pathogen *Listeria monocytogenes* disseminates through host tissues using a process called cell-to-cell spread. In this study, Sanderlin *et al*., discover that host proteins that normally regulate membrane trafficking and membrane remodeling in uninfected settings are also co-opted by *Listeria* to promote spread.

## Introduction

*Listeria monocytogenes* is a Gram-positive food-borne bacterium that causes listeriosis in humans (Radoshevich and Cossart, 2018). In some patients, complications such as meningitis or spontaneous abortions can occur because of this pathogen’s ability to infect many different cell types. A key to its pathogenesis is its ability to invade host cells and disseminate through tissues using a process called cell-to-cell spread. Cell-to-cell spread is characterized as a vesicular-mediated form of trafficking between host cells that allows *L. monocytogenes* to maintain access to cytosolic nutrients while also hiding from the humoral immune response (Lamason and Welch, 2016; Weddle and Agaisse, 2018). While much of the invasion process has been explored extensively (Radoshevich and Cossart, 2018), less is known about the molecular details of cell-to-cell spread.

To initiate spread, cytosolic *L. monocytogenes* first hijacks the host actin cytoskeleton to polymerize actin tails on the bacterial surface to promote cytosolic motility (Tilney and Portnoy, 1989; Welch et al., 1997). Once motile, it travels to the cell-cell junction and pushes a double-membrane protrusion from the donor cell into the recipient cell. This protrusion is eventually engulfed into a double-membrane vacuole, through unknown mechanisms, followed by rupture of the vacuole and bacterial escape into the recipient cell cytosol (Tilney and Portnoy, 1989; Robbins et al., 1999; Lamason et al., 2016). Early work suggested that spreading bacteria co-opted host pathways for efficient protrusion engulfment (Monack and Theriot, 2001); however, the identity of these factors has not been thoroughly evaluated.

Bacterial cell-to-cell spread resembles processes that occur in uninfected cells, in which adjacent cells exchange cytoplasm-containing vesicular compartments with their neighbors via trans-endocytosis. The details of this type of intercellular communication are still largely unknown, but factors important for cell-cell adhesion (e.g. cadherin), signaling (e.g. Eph receptors), endocytosis (e.g. clathrin), and exocytosis (e.g. endosomal sorting complexes required for transport (ESCRT) machinery) have been implicated in this process (Marston et al., 2003; Piehl et al., 2007; Sakurai et al., 2014; Gong et al., 2016). Because bacterial cell-to-cell spread mimics trans-endocytosis, we hypothesized that *L. monocytogenes* hijacks host proteins required for cell-cell adhesion, membrane trafficking, and membrane remodeling to promote spread.

In this study, we tested this hypothesis by conducting an RNAi screen that targeted 115 host genes and compared the requirements for these factors during *L. monocytogenes* spread. We discovered that 22 host genes are important for cell-to-cell spread. We further showed that loss of the endocytic proteins caveolin 1 and caveolin 2 or the membrane sculpting F-BAR protein PACSIN2 specifically impaired *L. monocytogenes* protrusion engulfment. We also observed localization of PACSIN2 to *L. monocytogenes* protrusions and defined a specific requirement for PACSIN2 in the recipient cell. Overall, our study shows that trafficking and membrane remodeling pathways are required for efficient bacterial spread, and reveals PACSIN2 as a key molecular player in this process. Our approach also highlights how investigating the mechanisms of bacterial spread may reveal fundamental insights into the regulation of host intercellular communication.

## Results and discussion

### RNAi screen reveals host genes that regulate bacterial cell-to-cell spread

Eukaryotic cells communicate with their neighbors and exchange material using intercellular communication processes such as trans-endocytosis. The molecular details are not well known, but what is clear is that trans-endocytosis allows cells to engulf membrane-bound cytoplasmic material from their neighbors (Rechavi et al., 2009). *L. monocytogenes* spreads through host cells by inducing double-membrane protrusions that must be engulfed by the recipient cell (Lamason and Welch, 2016; Weddle and Agaisse, 2018). Therefore, we predicted that pathways of intercellular communication, like trans-endocytosis, are hijacked to promote bacterial spread. To test this, we performed an RNAi screen targeting 115 genes implicated in cell-cell adhesion, membrane remodeling/curvature, and endocytosis. A549 cells were selected as the screening platform because this cell type is efficiently transfected, can form flat monolayers, and is easily infected by *L. monocytogenes*. Each screening plate contained ten replicates of a negative control siRNA (non-target, NT) and a positive control siRNA (Fig. 1, A and D). An siRNA targeting *ARPC4* (a subunit of the Arp2/3 complex) was selected as a positive control (Fig. 1 D) because silencing its expression impairs actin-based motility and spread (Chong et al., 2009; Talman et al., 2014). We also compared the RNAi-mediated spread defect to the *L. monocytogenes* Δ*actA* mutant, which is completely unable to spread due to a loss of actin-based motility (Kocks et al., 1992; Domann et al., 1992) (Fig. 1 D).

**Figure 1:**
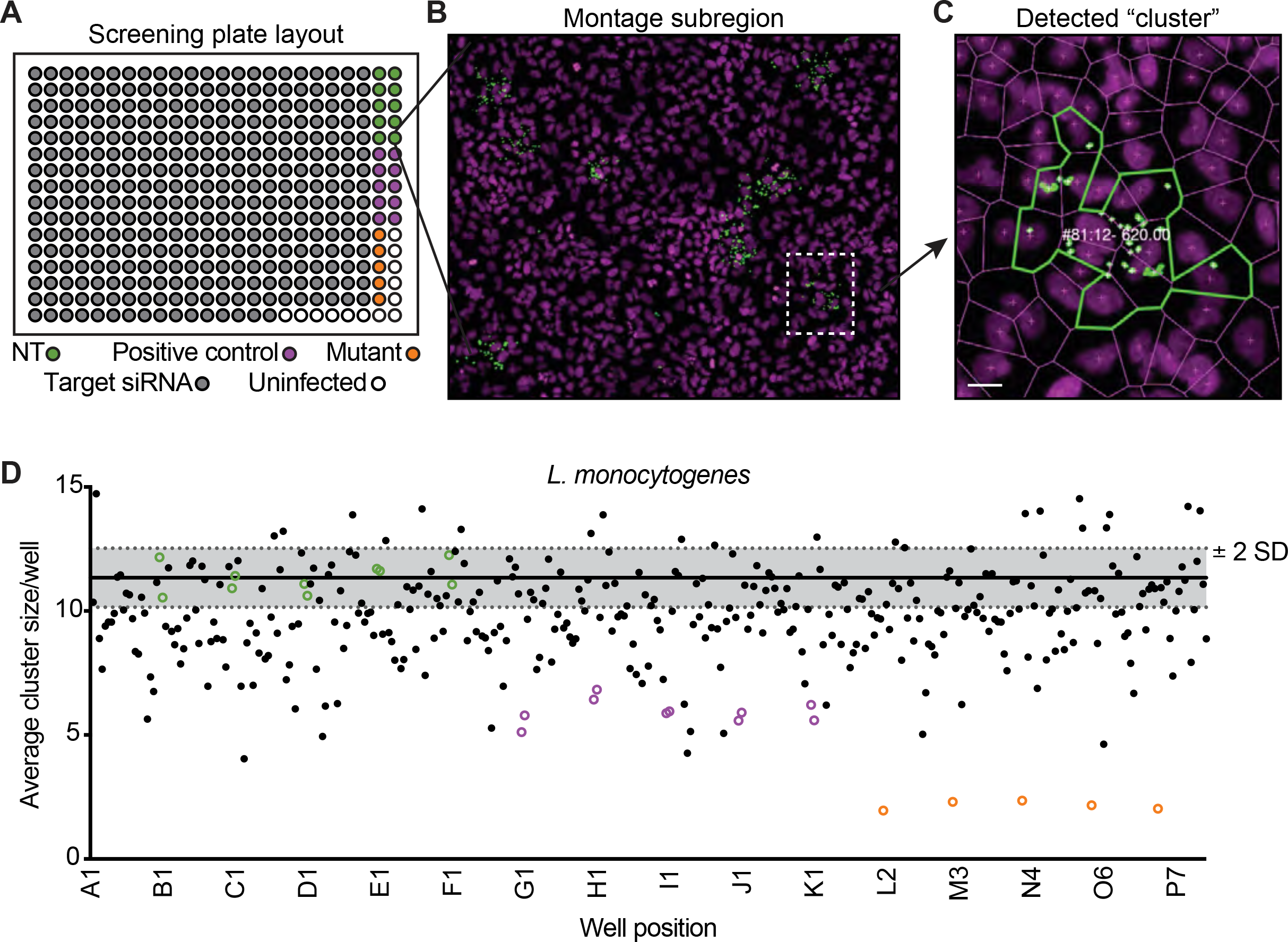
Primary RNAi screen reveals host genes that alter bacterial spread. (A) Screen plate layout with target siRNAs (grey circles), non-target siRNAs (NT, green circles), positive control siRNAs (purple circles), or bacterial spread mutants (orange circles). (B) A region of the full montage from the screen is shown with bacteria (green) and host nuclei (magenta). Region of interest (white dotted line) expanded in (C). (C) Example of a detected cluster (infectious focus) from the screen with the numbers representing “cluster #: infected cells - cumulative GFP signal.” Scale bar, 5 *µ*m. (D) Example run from the primary RNAi screen. NT (green circles), positive controls (purple circles), and mutant bacteria samples (orange circles) are shown along with screen siRNAs (black dots). Grey shaded regions represents ± 2 SD from the average cluster size (solid black line) of the ten NT control wells.

To measure spread efficiency, A549 cells were infected with a low multiplicity of infection (MOI) of GFP-expressing bacteria (LmGFP), and after 1 h of invasion, extracellular bacteria were washed away and killed with gentamicin to allow foci of infection to form. Infected samples were then fixed and stained to detect host nuclei and bacteria. An image analysis pipeline was used to quantify the number of infected cells per focus, as well as the number of foci per well (Fig. 1, B and C and Table S1). To measure spread efficiency in each well, we averaged the number of infected cells per focus and plotted those values for each screening run (Fig. 1 D and Table S1). Primary screen hits were selected if at least two siRNAs/gene showed a ± 2 standard deviation (SD) effect, relative to the negative control, across two biological replicates (Table S1). Based on this analysis, we found that silencing 29 genes reduced *L. monocytogenes* focus size and only one (*VPS24*) increased focus size.

We next determined whether any of these identified genes altered focus size indirectly by inhibiting actin-based motility or disrupting host cell monolayer integrity. RNAi-treated A549 cells were infected with TagBFP-expressing *L. monocytogenes* (LmTagBFP), and fixed samples were stained with phalloidin to detect F-actin. Images were collected with a confocal high-content imaging system to quantify the percentage of bacteria with actin tails (Fig. S1) and the extent of monolayer confluency (Fig. S2). Using this secondary screening approach, we found that RNAi-mediated silencing of *CAPZB* (F-actin capping protein) and *PVRL2* (cell adhesion protein Nectin 2) reduced actin tail frequencies (Fig. S1, B and C, and Table 1). CAPZB is a known regulator of *L. monocytogenes* actin-based motility (Loisel et al., 1999), while Nectin 2 has not been implicated and may indirectly affect this process. We also found that RNAi-mediated silencing of cell adhesion genes (e.g. *PVRL2*, *TLN1*, *JAM3*, and *CTNNA2*), membrane curvature/remodeling genes (e.g. *PSTPIP1* and *VPS28*), and *CDC42* (Rho GTPase) reduced cell confluency (Fig. S2, C and D and Table 1). In the end, we were able to remove eight genes with indirect effects, leaving 22 host genes that were important for bacterial spread (Table 1). This list included many cell-cell adhesion proteins, such as ZO-1 or E-cadherin, and several BAR domain family members, which sense or regulate host membrane curvature (Carman and Dominguez, 2018). Interestingly, we found that *L. monocytogenes* spread requires CAV1 and CAV2, which are core components of caveolin-mediated endocytosis in host cells (Busija et al., 2017) and may promote trafficking of the *L. monocytogenes* protrusion into the recipient cell. Therefore, we next examined whether caveolins regulated a specific step of *L. monocytogenes* spread.

**Table 1:**
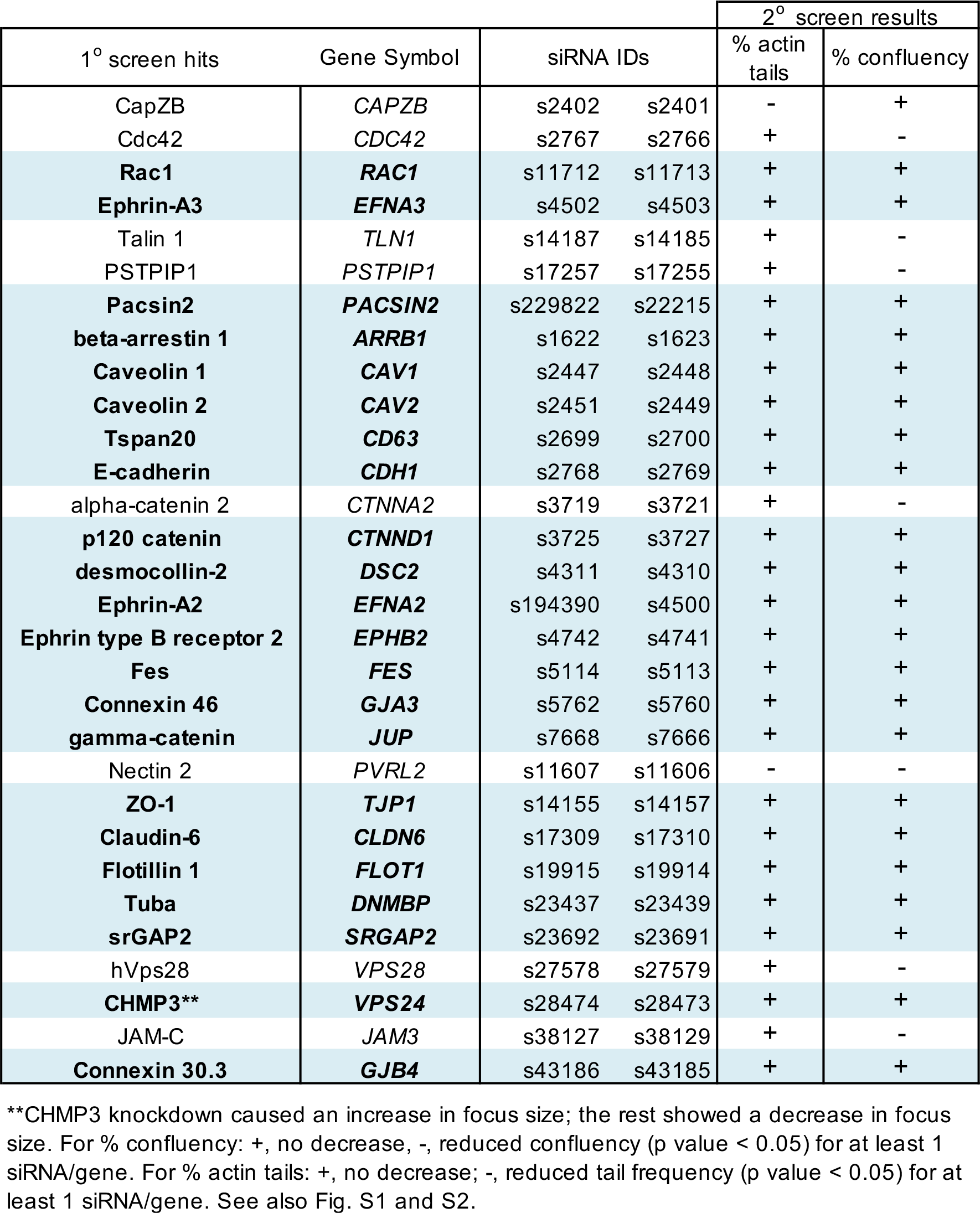
Combined results from the primary and secondary RNAi screens

|  |  |  |  | 2° screen results |  |
| --- | --- | --- | --- | --- | --- |
| 1° screen hits | Gene Symbol | siRNA IDs |  | % actin tails | % confluency |
| CapZB | *CAPZB* | s2402 | s2401 | - | + |
| Cdc42 | *CDC42* | s2767 | s2766 | + | - |
| **Rac1** | ***RAC1*** | s11712 | s11713 | + | + |
| **Ephrin-A3** | ***EFNA3*** | s4502 | s4503 | + | + |
| Talin 1 | *TLN1* | s14187 | s14185 | + | - |
| PSTPIP1 | *PSTPIP1* | s17257 | s17255 | + | - |
| **Pacsin2** | ***PACSIN2*** | s229822 | s22215 | + | + |
| **beta-arrestin 1** | ***ARRB1*** | s1622 | s1623 | + | + |
| **Caveolin 1** | ***CAV1*** | s2447 | s2448 | + | + |
| **Caveolin 2** | ***CAV2*** | s2451 | s2449 | + | + |
| **Tspan20** | ***CD63*** | s2699 | s2700 | + | + |
| **E-cadherin** | ***CDH1*** | s2768 | s2769 | + | + |
| alpha-catenin 2 | *CTNNA2* | s3719 | s3721 | + | - |
| **p120 catenin** | ***CTNND1*** | s3725 | s3727 | + | + |
| **desmocollin-2** | ***DSC2*** | s4311 | s4310 | + | + |
| **Ephrin-A2** | ***EFNA2*** | s194390 | s4500 | + | + |
| **Ephrin type B receptor 2** | ***EPHB2*** | s4742 | s4741 | + | + |
| **Fes** | ***FES*** | s5114 | s5113 | + | + |
| **Connexin 46** | ***GJA3*** | s5762 | s5760 | + | + |
| **gamma-catenin** | ***JUP*** | s7668 | s7666 | + | + |
| Nectin 2 | *PVRL2* | s11607 | s11606 | - | - |
| **ZO-1** | ***TJP1*** | s14155 | s14157 | + | + |
| **Claudin-6** | ***CLDN6*** | s17309 | s17310 | + | + |
| **Flotillin 1** | ***FLOT1*** | s19915 | s19914 | + | + |
| **Tuba** | ***DNMBP*** | s23437 | s23439 | + | + |
| **srGAP2** | ***SRGAP2*** | s23692 | s23691 | + | + |
| hVps28 | *VPS28* | s27578 | s27579 | + | - |
| **CHMP3\*\*** | ***VPS24*** | s28474 | s28473 | + | + |
| JAM-C | *JAM3* | s38127 | s38129 | + | - |
| **Connexin 30.3** | ***GJB4*** | s43186 | s43185 | + | + |

\*\*CHMP3 knockdown caused an increase in focus size; the rest showed a decrease in focus size. For % confluency: +, no decrease, -, reduced confluency (p value < 0.05) for at least 1 siRNA/gene. For % actin tails: +, no decrease; -, reduced tail frequency (p value < 0.05) for at least 1 siRNA/gene. See also Fig. S1 and S2.

### CAV1 and CAV2 promote protrusion engulfment during *L. monocytogenes* cell-to-cell spread

Endocytic proteins such as clathrin and caveolin regulate vesicle trafficking in host cells (Doherty and McMahon, 2009). Since bacterial spread resembles a vesicular form of trafficking, we predicted that caveolins might regulate *L. monocytogenes* spread by promoting protrusion engulfment into the neighboring cell. To test this, we infected A549 cells LmGFP after RNAi treatment and measured the percentage of bacteria in protrusions and vesicles (Fig. 2 A). As expected, we found that loss of *CAV1* or *CAV2* expression (Fig. 2 B) significantly increased the frequency of bacteria in protrusions (Fig. 2 C), suggesting that infectious size decreased because more bacteria were getting stuck in protrusions. We did not observe an apparent effect on bacteria-containing vesicle frequency (Fig. 2 D), possibly due to the low incidence of this phenotype. Altogether, our data suggest that loss of *CAV1* or *CAV2* expression reduces cell-to-cell spread by impairing protrusion engulfment efficiency.

**Figure 2:**
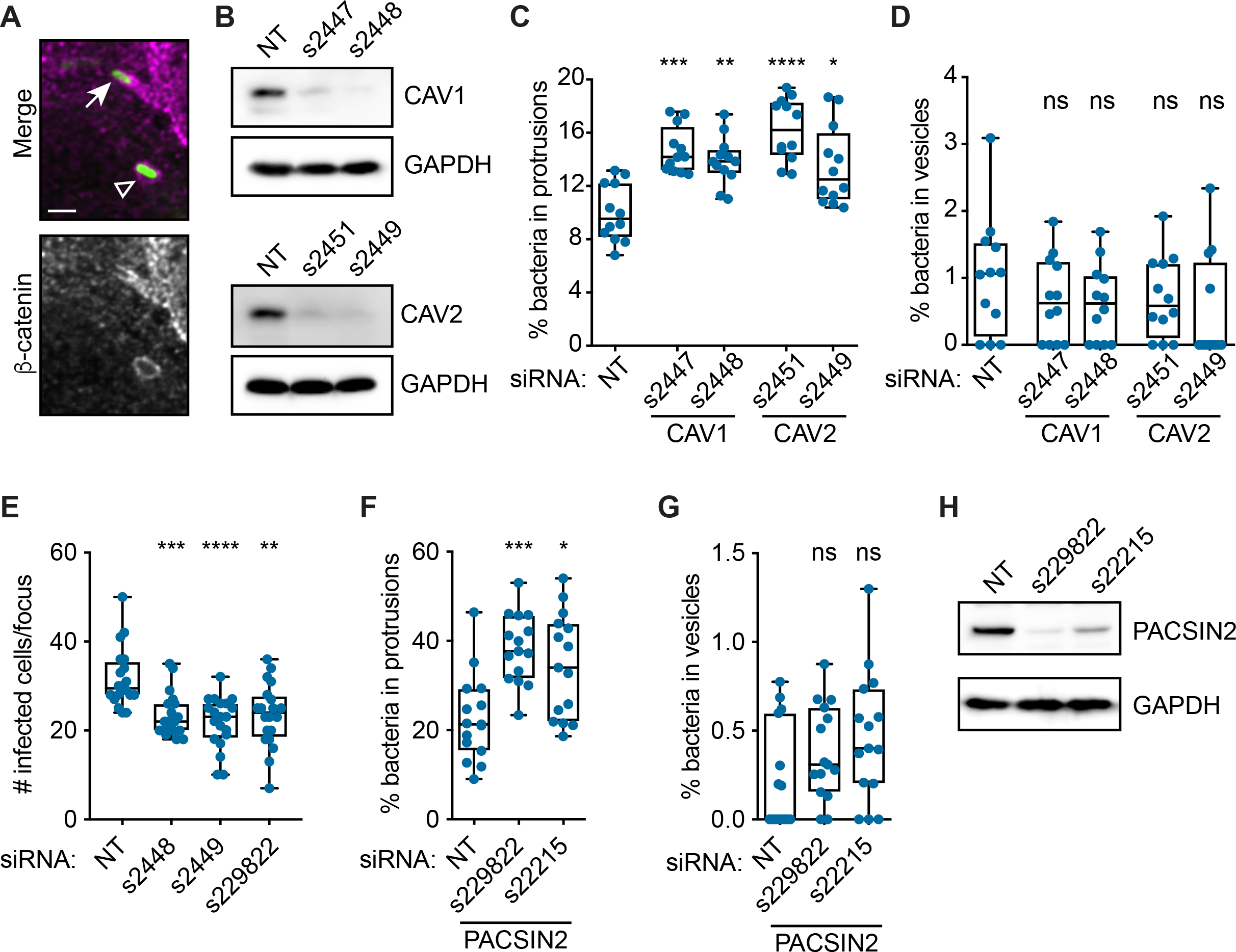
CAV1, CAV2, and PACSIN2 promote *L. monocytogenes* protrusion engulfment. (A) Image of infected A549 cells showing an example of LmGFP (green) in either a protrusion (arrow) or a vesicle (open triangle). Host membrane detected via β-catenin immunofluorescence (grey; magenta in merge). Scale bar, 2 *µ*m. (B) Western blot of CAV1, CAV2, and GAPDH (loading control) expression in A549 cells after indicated RNAi treatments. (C, D) Percentage of *L. monocytogenes* in protrusions (C) or vesicles (D) in A549 cells after RNAi-silencing of *CAV1* or *CAV2*. (E) Infectious focus assay results after RNAi-mediated silencing of *CAV1* (s2448), *CAV2* (s2449), or *PACSIN2* (s229822) expression in Caco-2 BBe1 cells. (F, G) Percentage of *L. monocytogenes* in protrusions (F) or vesicles (G) in A549 cells after RNAi-silencing of *PACSIN2*. (H) Western blot of PACSIN2 and GAPDH (loading control) expression in A549 cells after indicated RNAi treatments. Significance determined relative to the NT siRNA control using a Kruskal-Wallis ANOVA with Dunn’s multiple comparison test. *p < 0.05, **p < 0.01, ***p < 0.001, ****p < 0.0001. ns, not significant.

We also investigated whether CAV1 and CAV2 promoted spread in intestinal cells, a physiological target of *L. monocytogenes* (Radoshevich and Cossart, 2018). We treated Caco-2 BBe1 (human enterocytic) cells with siRNAs against *CAV1* and *CAV2* and discovered that silencing the expression of *CAV1* and *CAV2* significantly reduced infectious focus size in Caco-2 BBe1 cells (Fig. 2 E). This suggests that CAV1 and CAV2 promote spread in multiple cell types. Given the importance of caveolins to *L. monocytogenes* spread, we next asked whether other caveolin regulators from our screen promoted protrusion engulfment.

### The F-BAR protein PACSIN2 promotes protrusion engulfment during *L. monocytogenes* cell-to-cell spread

Caveolae biogenesis is coordinated by multiple proteins acting in concert with caveolins to sculpt the membrane (Echarri and Del Pozo, 2015; Parton et al., 2018). The F-BAR protein PACSIN2 (aka syndapin-II) is one such component that binds to CAV1 and regulates caveolae biogenesis and endocytosis (Senju et al., 2011; Hansen et al., 2011). Intriguingly, our RNAi screen revealed that PACSIN2 might also promote bacterial cell-to-cell spread (Table 1). Therefore, we tested whether PACSIN2 also promoted *L. monocytogenes* protrusion engulfment. We examined protrusion and vesicle frequency in A549 cells as above and found that silencing *PACSIN2* expression significantly increased the percentage of bacteria in protrusions (Fig. 2, F and H), but did not affect vesicle frequency (Fig. 2 G). We also measured *L. monocytogenes* spread in Caco-2 BBe1 cells and found that RNAi-mediated silencing of *PACSIN2* expression reduced infectious foci size (Fig. 2 D). Overall, these results suggest that, just like CAV1 and CAV2, PACSIN2 promotes spread in multiple cell types and acts by promoting protrusion engulfment.

### PACSIN2 localizes to *L. monocytogenes*-containing protrusions and is required in the recipient cell to promote spread

Since caveolins and PACSIN2 regulate protrusion engulfment, we wanted to see if they were localized to the protrusion membrane. We first examined the localization of endogenous CAV1 to *L. monocytogenes*-containing protrusions since overexpression of caveolins leads to aberrant structures (Parton and Del Pozo, 2013). Infected A549 cells were fixed and stained for endogenous caveolins, and the percentage of bacterial protrusions with caveolin localization was counted. CAV1 was observed in puncta around the plasma membrane, as had been previously shown in uninfected cells (Hansen et al., 2011). Interestingly, we only observed CAV1 localization on 1.7% (SD 1.3%) of the LmTagBFP-containing protrusions across two biological replicates (from a total of 2232 protrusions). This pattern of CAV1 localization was also restricted to shorter protrusions (< 3 *µ*m) near the cell-cell junction (Fig. 3 A). This low frequency of colocalization may not be surprising since, relative to adipocytes or muscle cells, epithelial cells do not contain abundant caveolae (Parton and Del Pozo, 2013; Parton et al., 2018). We were also unable to detect dynamic recruitment of a CAV1-mNeonGreen fusion in gene-edited A549 cells using live cell microscopy (data not shown). Therefore, we speculate that if caveolins are acting at the protrusion membrane, they may be at very low levels or are stably maintained in the membrane (Tagawa et al., 2005; Parton and Del Pozo, 2013) to recruit other factors, such as PACSIN2.

**Figure 3:**
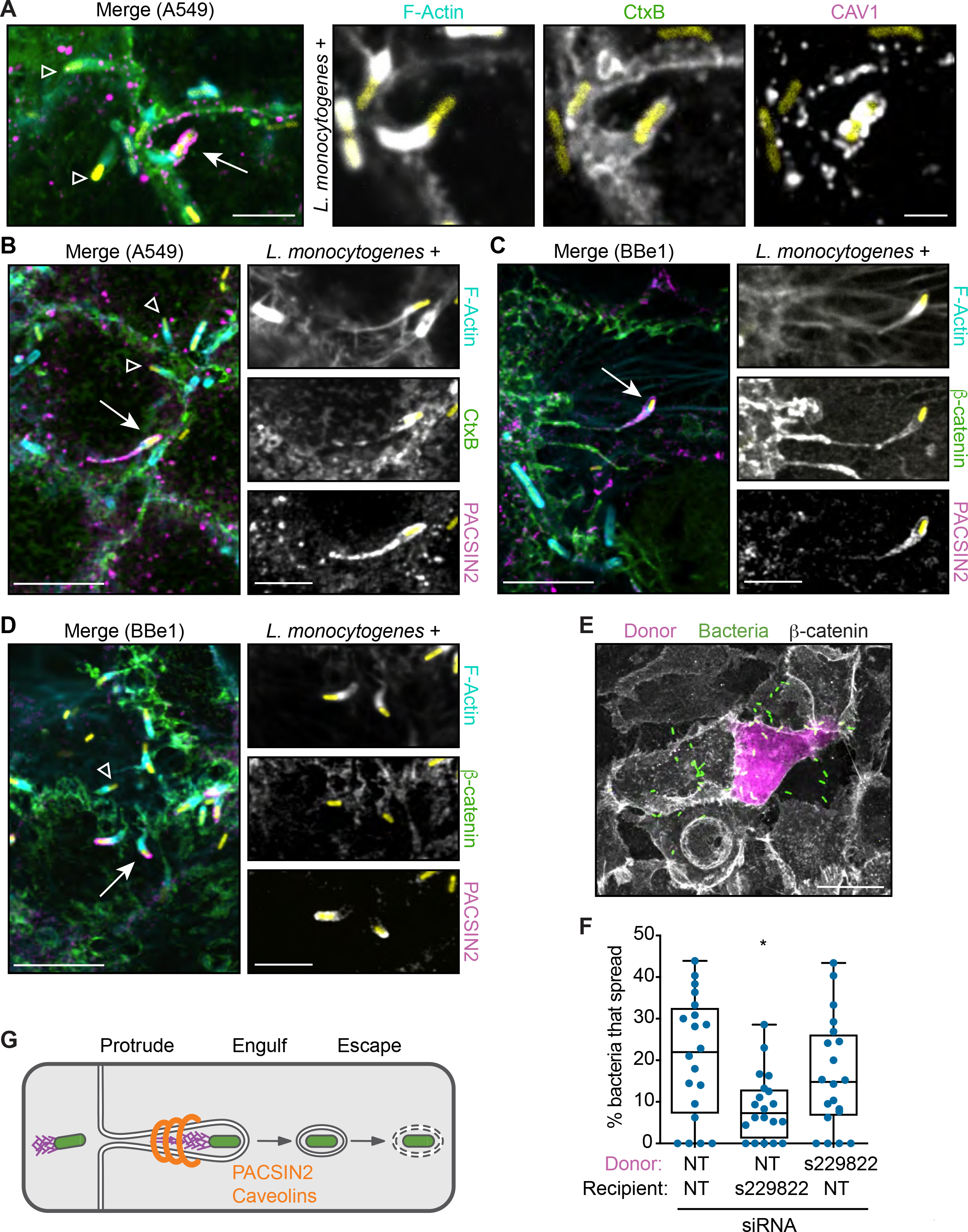
PACSIN2 is required in the recipient cell and localizes to *L. monocytogenes* protrusions. (A) Localization of endogenous CAV1 (magenta) in A549 cells infected with LmTagBFP (yellow). Scale bar, 5 *µ*m (inset, 2 *µ*m). (B-D) Localization of endogenous PACSIN2 (magenta) in A549 (B) or Caco-2 BBe1 (C, D) cells after infection with LmTagBFP (yellow). Scale bar, 10 *µ*m (inset, 5 *µ*m). For (A-D), membranes (green) were detected with CtxB (A, B) or an antibody to β-catenin (C, D) and F-actin was labeled with phalloidin (cyan). Arrow indicates host factor-positive protrusion used in inset; open triangle indicates host factor-negative protrusions. (E) Image from the mixed cell infectious focus assay, indicating the donor cell (magenta), LmGFP (green), and host membrane detected via β-catenin immunofluorescence (grey). Scale bar, 20 *µ*m. (F) Mixed cell infectious focus assay results in A549 cells after silencing *PACSIN2* expression. (G) Diagram of *L. monocytogenes* cell-to-cell spread indicating when PACSIN2 and caveolins promote spread.

PACSIN2 is recruited to caveolae via its F-BAR domain (Hansen et al., 2011) and it directly interacts with CAV1 (Senju et al., 2011). This crescent-shaped F-BAR module also binds to positively curved membranes and can induce membrane tubulation when overexpressed in cells (Senju et al., 2011). Therefore, we predicted that PACSIN2 would localize to protrusions. To test this, cells were infected with LmTagBFP and endogenous PACSIN2 protein detected by immunofluorescence. As previously published, PACSIN2 localized to small puncta and along tubules near the plasma membrane (Hansen et al., 2011; Dorland et al., 2016). More importantly, we observed colocalization of PACSIN2 with approximately 18.9% (SD 6.2%) of the LmTagBFP-containing protrusions in A549 cells and 7.9% (SD 4.3%) of the LmTagBFP-containing protrusions in Caco-2 BBe1 cells. PACSIN2 was often found along the length of long protrusions with skinny membranous stalks and a collapsed actin network (Fig. 3, B and C), suggesting that PACSIN2 accumulated on late-stage protrusions. A subset of the PACSIN2-containing protrusions in Caco-2 BBe1 (13% ± 13.6%) were shorter and exhibited specific localization of PACSIN2 to the tips of the protrusions (Fig. 3 D), suggesting PACSIN2 may assemble on an emerging protrusion and accumulate during protrusion elongation.

Given the striking localization of PACSIN2, we also predicted that it specifically acted in the recipient cell to facilitate protrusion engulfment. To test this, we performed a mixed-cell infectious focus assay to determine the requirement for PACSIN2 in donor or recipient host cells. In this assay, donor host cells expressing a red fluorophore (TagRFP-T) are pre-infected with LmGFP and then mixed with unlabeled recipient cells (Fig. 3 E). Since each population is separate before the mix, we can independently modulate the expression levels of our host targets in each host cell population via RNAi-mediated silencing. Using this assay, we found that spread was only reduced when *PACSIN2* expression was silenced in the recipient cells (Fig. 3 F), suggesting that PACSIN2 acts specifically in the recipient cell. Taken together with the work above, our data support a model whereby caveolins and the F-BAR protein PACSIN2 promote *L. monocytogenes* cell-to-cell spread by regulating protrusion engulfment into the recipient cell (Fig. 3 G).

## Conclusion

The direct exchange of organelles and cytoplasmic material between cells allows multicellular organisms to communicate and execute essential functions (Rechavi et al., 2009; Mittelbrunn and Sánchez-Madrid, 2012; Murray and Krasnodembskaya, 2019). Although the details are not well understood, we hypothesized that specific proteins would be required (e.g. cell-cell adhesion, membrane remodeling, and membrane trafficking) and that they would be co-opted by spreading bacteria. By focusing on this process of host intercellular communication, we identified new regulators of *L. monocytogenes* cell-to-cell spread. One key finding was that host membrane curvature and trafficking proteins (e.g. PACSIN2 and caveolins) promoted protrusion engulfment during *L. monocytogenes* spread. Although our data do not reveal their mechanisms of action, our work highlights how exploring these questions could provide new insights into the types of host processes co-opted by spreading bacteria and the regulation of host intercellular communication in uninfected settings.

In addition to exploring the mechanisms of PACSIN2 and caveolins, it will be essential to reveal how other hits from our screen are co-opted for *L. monocytogenes* spread. Nearly all of the hits appear to promote spread, and we predict that they act by regulating protrusion initiation or engulfment. Interestingly, only one gene (*VPS24*) increased infectious focus size when its expression was silenced, suggesting it normally limits spread. *VPS24* encodes CHMP3, which is a component of the ESCRT-III machinery that serves to constrict and sever membranes (Christ et al., 2017). Precisely how the loss of CHMP3 enhances spread is not clear, but previous work has shown that the ESCRT machinery can be hijacked by other pathogens such as retroviruses (Scourfield and Martin-Serrano, 2017).

Finally, it will be important to examine whether similar or different host processes are required for other spreading bacteria. Intriguingly, *Shigella flexneri* uses a non-canonical form of clathrin-mediated endocytosis to spread (Fukumatsu et al., 2012). Our data show that the clathrin machinery is not required for *L. monocytogenes* spread, suggesting different pathogenic strategies have evolved. Thus, examining the species-specific mechanisms of cell-to-cell spread may uncover how different host processes are co-opted by bacterial pathogens and how this impacts their modes of pathogenesis.

---

**Figure S1: Secondary RNAi screen reveals host genes that affect actin-based motility.** (A) Example images of phalloidin-stained samples used to measure the frequency of TagBFP-expressing *L. monocytogenes* with actin tails. Arrows indicate bacteria (magenta) with actin tails (green). Scale bar, 10 *µ*m (inset, 5 *µ*m). (B) Pooled data from two biological replicates of the screens are shown. Each dot represents the average frequency of bacteria with actin tails per well (3 wells/siRNA were done in each run of the screen). Significance determined relative to the NT siRNA control using a Kruskal-Wallis ANOVA with an uncorrected Dunn’s test. Note that for visibility, the data in (B) are split into two graphs and the NT data duplicated. *p < 0.05, **p < 0.01, ***p < 0.001, ****p < 0.0001.

**Figure S2: Secondary RNAi screen reveals host genes reduce host monolayer integrity.** (A, B) Phalloidin-stained samples were imaged (left) and then a binary mask was applied (right) to quantify the percentage of surface area covered by cells (i.e. % confluency) from intact (A) and disrupted (B) monolayers. Scale bar, 10 *µ*m. (C) Pooled data from two biological replicates of the screens are shown. Each dot represents the average percentage of confluency per well (3 wells/siRNA were done in each run of the screen). Significance determined relative to the NT siRNA control using a Kruskal-Wallis ANOVA with an uncorrected Dunn’s test. Note that for visibility, the data in (C) are split into two graphs and the NT data duplicated. *p < 0.05, **p < 0.01, ***p < 0.001, ****p < 0.0001.

**Table S1: Primary siRNA screen results**

**Table S2: List of siRNAs used in the RNAi screens**

## Acknowledgments

First, we would like to acknowledge the incredible generosity of Matthew Welch. This work was started in the Welch lab while R.L.L was a postdoc, and Matt kindly allowed R.L.L to complete the study in her own lab. This is a rare gift from a truly special mentor. We would also like to thank Jon McGinn, Adam Martin, and Matthew Welch for their critical reading of the manuscript. We thank Erin Benanti, Michelle Reniere, and Dan Portnoy for sharing reagents. Invaluable technical help for the RNAi screen was provided by Trish Birk, Mary West, Pingping He, and Andreas Ettinger. This work was performed in part at the High Throughput Screening Facility (UCB) and the Keck Imaging Facility (Whitehead/MIT). This work was supported in part by the NIH Pre-Doctoral Training Grants T32GM007287. R.L.L is supported by NIH grant R00GM115765.

## Author Contributions

Conceptualization and Methodology, Rebecca Lamason, Indro Fedrigo, and Vida Ahyong; Software, Indro Fedrigo, Vida Ahyong, and Rebecca Lamason; Investigation, Allen Sanderlin, Cassandra Vondrak, Arianna Scricco, and Rebecca Lamason; Writing, Allen Sanderlin, Cassandra Vondrak, and Rebecca Lamason; Supervision, Rebecca Lamason.

## Materials and methods

### Cell lines

A549 cells were obtained from the University of California, Berkeley tissue culture facility. Caco-2 BBe1 cells were kindly provided by the laboratory of Dr. Marcia Goldberg (Massachusetts General Hospital). All cells were grown at 37°C in 5% CO_2_ and maintained in DMEM (Invitrogen) containing 10% fetal bovine serum (FBS; Atlas Biologicals). A549 cells stably expressing TagRFP-T (A549-TRT) were generated using retroviral transduction, as previously described (Lamason et al., 2016).

### Bacterial Strains

GFP-expressing wild-type *L. monocytogenes* (Shen and Higgins, 2005) were kindly provided by Dr. Daniel Portnoy (UC Berkeley). The GFP-expressing Δ*actA L. monocytogenes* strain was a gift from Drs. Michelle Reniere and Dan Portnoy and was generated by integrating pPL2-gfp (Lauer et al., 2002) into the DP-L3078 (Δ*actA*) strain (Skoble et al., 2000). The TagBFP-expressing *L. monocytogenes* strain (LmTagBFP, strain PL 1949) was a kind gift from Dr. Erin Benanti (Aduro Biotech). To generate LmTagBFP, TagBFP was codon optimized for *L. monocytogenes* expression by ATUM (Newark, CA) and cloned downstream of the *actA* promoter in pPL2. The TagBFP expression cassette was integrated at the tRNA-Arg locus of *L. monocytogenes* by site-specific integration (Lauer et al., 2002).

### Primary siRNA Screen

A Silencer Select siRNA custom library was purchased from Ambion (Life Technologies) (see Table S2). The master library plate was made by diluting siRNAs to 0.25 *µ*M and arraying them into a deepwell 384-well storage microplate (Corning Axygen P-384-120SQ-C) as indicated in Fig 1A. A non-target siRNA (Negative control #1 siRNA, #4390843) was used as a negative control and an siRNA against the *ARPC4* gene (Ambion AM16708) was selected as a positive control. Note, this reagent is from Ambion’s Silencer collection, which requires more siRNA per reaction. To set-up screen plates, 1 *µ*l of siRNA from each well of the master plate was spotted onto a 384-well *µ*Clear black plate (Greiner 781091) using an Agilent Velocity 11 Bravo liquid handler. A549 cells were then reverse transfected in these plates via Lipofectamine RNAiMAX using a Multidrop Combi Dispenser (Thermo Scientific) to dispense cells and reagents, which resulted in a final siRNA concentration of 5 nM (or 50 nM for the *ARPC4*-specific siRNA) and 4.4 × 10^3^ cells per well.

To measure *L. monocytogenes* spread efficiency via the infectious focus assay, transfected cells were infected 72 h post-transfection at a multiplicity of infection (MOI) of 0.08 and plates were centrifuged at 200 × g for 5 min at 25°C and incubated at 37°C for 1 h. Samples were washed 1 time with PBS before adding complete media with 10 *µ*g/ml gentamicin, and the infection progressed for 4.5 h at 37°C until fixation and staining. Infected cells were fixed in 4% PFA in 1× phosphate buffered saline (PBS) for 10 min at room temperature and washed twice in 1× PBS. Staining was then performed on the Agilent Velocity 11 Bravo liquid handler as follows: fixed cells were permeabilized with 0.5% Triton X-100 (T×100) for 5 min at room temperature, washed once with 1× PBS and incubated with primary and secondary antibodies diluted with 2% bovine serum albumin (BSA) in 1× PBS. To increase the detection of our GFP-expressing bacteria, we found that staining for each strain was necessary. Therefore, *L. monocytogenes* stained using rabbit anti-*Listeria* (Difco 223021), and nuclei stained with Hoechst 33342 (Molecular Probes). Stained cells were covered with 50 *µ*l per well of sterile 50% glycerol in 1× PBS before imaging.

Primary screen plates were imaged on an ImageXpress Micro High Content Imaging System (Molecular Devices) using a 20× ELWD objective. In order to sample most of the well surface area, a 5×5 or 6×6 grid of images were collected and processed and stitched together using Fiji (ImageJ) (Schindelin et al., 2012) to create a montage. Montages were then analyzed using CellProfiler (Carpenter et al., 2006) to detect the positions of bacteria and nuclei in the stitched images and to estimate cell boundaries. This information was then analyzed using R (RStudio) to determine the number of infected cells per focus of infection (aka average cluster size) and the number of foci per well. Bacterial density was measured by quantifying the cumulative GFP signal intensity per focus and the values representing ≤ 4 bacteria determined manually. Foci with less than 4 bacteria were excluded from the analysis since they likely represented an unproductive infection. For each well, the average number of infected cells per focus (or average cluster size) was calculated and this value was used in selecting hits.

To select hits, two independent runs of the screens were done on different days and the results were compared between these two runs. In each run, the standard deviation (SD) was calculated for the 10 replicates of the NT control samples and this was used to measure the extent of the effects of each of our test siRNAs. siRNAs that increased or decreased the average cluster size per well by at least 2 SD were considered to have an effect. In each plate, every target gene was represented by 3 different siRNAs, each within their own well. Therefore, we only deemed a “hit” when at least 2 siRNAs per gene showed a consistent 2 SD effect across both runs of the screen (Table S1).

### Secondary siRNA Screen

To conduct the secondary siRNA screens, two siRNAs per gene were selected from the primary screen based on their ability to generate a consistent spread phenotype. To examine the effects of these siRNAs on host cell confluency and actin tail frequency, 2 × 10^4^ A549 cells were reverse transfected with 5nM siRNA via Lipofectamine RNAiMAX in a 384-well *µ*Clear black plate (Greiner 781091). In each plate, each siRNA was set-up in triplicate and two full runs of the screen were completed for each infection. Infections with TagBFP-expressing *L. monocytogenes* (LmTagBFP) were started 4 d after transfection at an MOI of 0.25. After adding bacteria, infected cells were centrifuged at 200 × g for 5 min at 25°C and incubated at 37°C for 4 h. At the end of the incubations, samples were fixed in 4% PFA in 1× PBS for 10 min at room temperature and washed once in 1× PBS. Fixed cells were permeabilized with 0.5% T×100 for 5 min at room temperature, washed three times with 1× PBS, and incubated for 1 h at room temperature with Alexa Fluor 488 phalloidin (ThermoFisher) diluted in 2% bovine serum albumin (BSA) in 1× PBS. Samples were then washed 3 times in 1× PBS and fresh 1× PBS was used to overlay samples before imaging.

Imaging of screen plates was done on an OperaPhenix High Content Imaging System (Perkin Elmer) using a 63X water immersion objective. For each well, 4 images were collected from randomly selected regions, processed using Fiji (Schindelin et al., 2012), and analyzed using CellProfiler (Carpenter et al., 2006) to quantify the number of bacteria per field of view and the cell confluency. For the cell confluency measurement, images with phalloidin signal only were converted to binary images and the % of confluency was quantified as the fraction of the image surface area covered by cells. To calculate the actin tail frequency, each image was manually scored for the number of bacteria with tails at least 1 bug length long and this number was divided by the number of total bugs in the field. For every well, the average phenotype across the 4 separate images was calculated and these well-based averages were plotted in Fig. S 1 and 2.

### Infectious Focus Assay

To measure *L. monocytogenes* spread efficiency in Caco-2 BBe1, 12 mm sterile coverslips in a 24-well plate were first coated with 15 *µ*g collagen-I (Sigma C7661) for 1 h at room temperature, washed 3 times with 1× PBS, and stored dry at 4°C until use. Then, 2 × 10^5^ cells were plated onto the collagen-coated coverslips and incubated for 24 h at 37°C. The next day, the cells were transfected with 10 nM siRNA using Lipofectamine RNAiMAX (ThermoFisher). 72 h after transfection, samples were infected with LmGFP at an MOI of 0.01, spun down for 5 min at 200 × g at 25°C, and then incubated for 1 h at 37°C. Samples were then washed 3 times in 1× PBS before adding complete media with 10 *µ*g/ml gentamicin. Infections proceeded for an additional 9-10 h at 37°C followed by fixation in 4% PFA in 1× PBS for 10 min at room temperature. Fixed samples were washed once in 1× PBS and then incubated with 0.1 M glycine in 1× PBS for 10 min at room temperature. Samples were then washed 3 times with 1× PBS, permeabilized with 0.5% T×100 for 5 min at room temperature and washed once with 1× PBS. Samples were then blocked in blocking buffer (2% BSA in 1× PBS) and incubated with primary and secondary antibodies diluted in blocking buffer. Hoechst 33342 (Molecular Probes) was used to stain the nucleus, mouse anti-β-catenin (Cell Signaling 2677S) was used to detect the host plasma membrane, and Alexa Fluor 647 phalloidin (ThermoFisher) was used to detect F-actin. Images were captured on our Olympus IXplore Spin microscope system with a Yokogawa CSU-W1 spinning disc unit, 60X UPlanSApo (1.3 NA) and 100X UPlanSApo (1.35 NA) objectives, an ORCA-Flash4.0 sCMOS camera, and CellSens software. 15-20 infectious foci were imaged per condition, images were processed in Fiji (Schindelin et al., 2012) and the number of infected cells/focus calculated.

### Protrusion and vesicle frequency assay

To measure the percent of bacteria in protrusions and vesicles, 0.75 × 10^5^ A549 cells were plated on 12 mm coverslips in 24-well plates. 24 h later, cells were transfected with 5nM siRNA via Lipofectamine RNAiMAX (ThermoFisher). 72 h after transfection, cells were infected with LmGFP at an MOI of 1-1.3 by centrifuging plates for 5 min at 200 × g at 25°C and subsequently incubating at 37°C for 4.5 h before fixation with 4% PFA/1x PBS. Fixed samples were stained as above for the Caco-2 BBe1 cell infectious focus assay. Images were captured on either a Nikon Ti-E inverted microscope with an Andor Revolution spinning disc confocal, Andor Zyla 5.5 sCMOS camera, and a 100X PlanApo objective (Fig. 2 C and D) WM Keck Microscopy Facility, or our Olympus IXplore Spin microscope system described above with a 100X UPlanSApo (1.35 NA) objective (Fig. 2 F and G). 12-15 fields of view were captured across 2-3 coverslips per experiment (with each field containing ~100-1000 bacteria). Images were processed in Fiji (Schindelin et al., 2012) and the average percentage of bacteria in protrusions and vesicles was calculated. Note, we suspect that the differences in image resolution acquired on the two different systems account for the different frequencies seen in the NT controls from Fig. 2 C, D, F, and G. To confirm this, s2448 was used as a positive control when acquiring the PACSIN2 data (Fig. 2 F), and we were able to see an increase in protrusion frequency compared to the NT control (data not shown).

### Mixed cell infectious focus assay

To set-up the mixed cell spread assays, 5.4 × 10^3^ donor cells/well (A549-TRT) in 96-well plates (Nunc/ThermoFisher Scientific), and 2 × 10^5^ recipient cells/well (A549) in 6-well plates (Genesee Scientific) were reverse transfected with 5 nM siRNA using Lipofectamine RNAiMAX (ThermoFisher). 72 h after plating, donors were infected with 2 × 10^6^ colony forming units (cfu) of LmGFP and plates were centrifuged at 200 × g for 5 min at 25°C and incubated at 37°C for 30 min. All samples (donor and recipient cells) were then washed once with PBS, lifted with 37°C 1× citric saline (135 mM KCl, 15 mM sodium citrate), and recovered in complete media. Cells were pelleted and then washed twice in complete media to remove residual citric saline. Donor cell pellets were resuspended in 50 *µ*l and recipient cell pellets resuspended in 1.3 ml complete media + 10 *µ*g/ml gentamicin. Cells were then mixed at a cell ratio of 1:125 (5.3 *µ*l donors plus 500 *µ*l recipients) and plated onto 12 mm coverslips in a 24-well plate. To facilitate fast adherence to the glass, the coverslips were pre-coated overnight at 4°C with 5 *µ*g/ml fibronectin (EMD Millipore) in 1× PBS and washed with 1× PBS immediately before added the cell mixtures. Infections were allowed to progress at 37°C for 4-4.5 h until fixation and staining. Staining proceeded as above for the Caco-2 BBe1 cell infectious focus assay, except samples were only stained with mouse anti-β-catenin (Cell Signaling 2677S). To quantify spread, 20 individual foci were imaged on our Olympus IXplore Spin system. Images were then processed in Fiji (Schindelin et al., 2012) and the percentage of bacteria per focus that had spread to recipient cells was calculated.

### Immunolocalization of host factors

To measure the frequency of PACSIN2, CAV1, and CAV2 localization during *L. monocytogenes* cell-to-cell spread, 1.5 × 10^5^ A549 cells were first plated onto 12 mm coverslips in a 24-well plate. 72 h later, cells were infected with 1×10^5^ colony forming units (cfu) of LmTagBFP and plates were centrifuged at 200 × g for 5 min at 25°C. Plates were then placed at 37°C and infection proceeded for 5 h before fixation in 4% PFA in 1× PBS for 10 min at room temperature.

For localization studies in Caco-2 BBe1 cells, each coverslip was first coated with 15 *µ*g collagen-I (Sigma C7661) for 1 h at room temperature, washed 3 times with 1× PBS, and stored dry at 4°C until use. Then, 1.5 × 10^5^ Caco-2 BBe1 cells were added to coated coverslips and after 48 h, the media was changed. The next day, cells were infected with 2 × 10^6^ cfu of LmTagBFP and plates centrifuged at 200 × g for 5 min at 25°C. Infections in Caco-2 BBe1 cells progressed for 9.5 h at 37°C followed by fixation in 4% PFA in 1× PBS for 10 min at room temperature.

Once fixed, all samples were washed once in 1× PBS and then incubated with 0.1 M glycine in 1× PBS for 10 min at room temperature. Samples were then washed 3 times with 1× PBS, permeabilized with 0.5% T×100 for 5 min at room temperature and washed once with 1× PBS. Samples were then incubated with Image-iT FX signal enhancer (ThermoFisher) for 30 min at room temperature, washed once with 1× PBS, then blocked with blocking buffer (2% BSA in 1× PBS) for 30 min at room temperature. Primary and secondary antibodies were diluted in blocking buffer and incubated for 1 h each at room temperature. The following stains and antibodies were used: rabbit anti-PACSIN2 (Abgent AB8088b), rabbit anti-CAV1 (Abcam ab2910), mouse anti-CAV2 (BD 610684), Alexa Fluor 647 phalloidin (ThermoFisher), mouse anti-β-catenin (Cell Signaling 2677S), and Alexa Fluor 488 conjugated Cholera Toxin Subunit B (Invitrogen C34778). Ten random fields of view per sample were imaged on our Olympus IXplore Spin microscope system using a 100X UPlanSApo (1.35 NA) objective. Images were processed in Fiji (Schindelin et al., 2012) and the percentage of protrusions showing colocalization with the host proteins of interest was calculated across two independent experiments. Percentages were collected across two biological replicates in both A549 cells (from a total of 863 protrusions) and Caco-2 BBe1 cells (from a total of 1899 protrusions).

### Western Blot Analysis

To determine the extent of knockdown, cells were first lysed in immunoprecipitation (IP) lysis buffer (50 mM HEPES, 150 mM NaCl, 1 mM EDTA, 10% Glycerol, 1% Igepal) on ice for 10 min. The cell debris was then cleared via centrifugation at 16,100 × g 4°C for 10 min. Lysates were analyzed by Western blotting using rabbit anti-PACSIN2 (Agent AP8088b), mouse anti-CAV1 (BD 610406), mouse anti-CAV2 (BD 610684), rabbit anti-CAV1 (for use with Caco-2 BBe1 cells; Abcam ab2910), and mouse anti-GAPDH (AM4300, Ambion).

### Statistical Analysis

Statistical analysis was performed in GraphPad PRISM 8 and the parameters and significance are reported in the Figures and the Figure Legends. Asterisks denote statistical significance as: *, p < 0.05; **, p < 0.01; ***, p < 0.001; ****, p < 0.0001, compared to indicated controls. For graphs depicted as box plots, boxes outline the 25th and 75th percentiles, midlines denote medians, and whiskers show the minimum and maximum values.

