## Supplemental Figures and Tables for "RNAi screen reveals a role for PACSIN2 and caveolins during bacterial cell-to-cell spread"

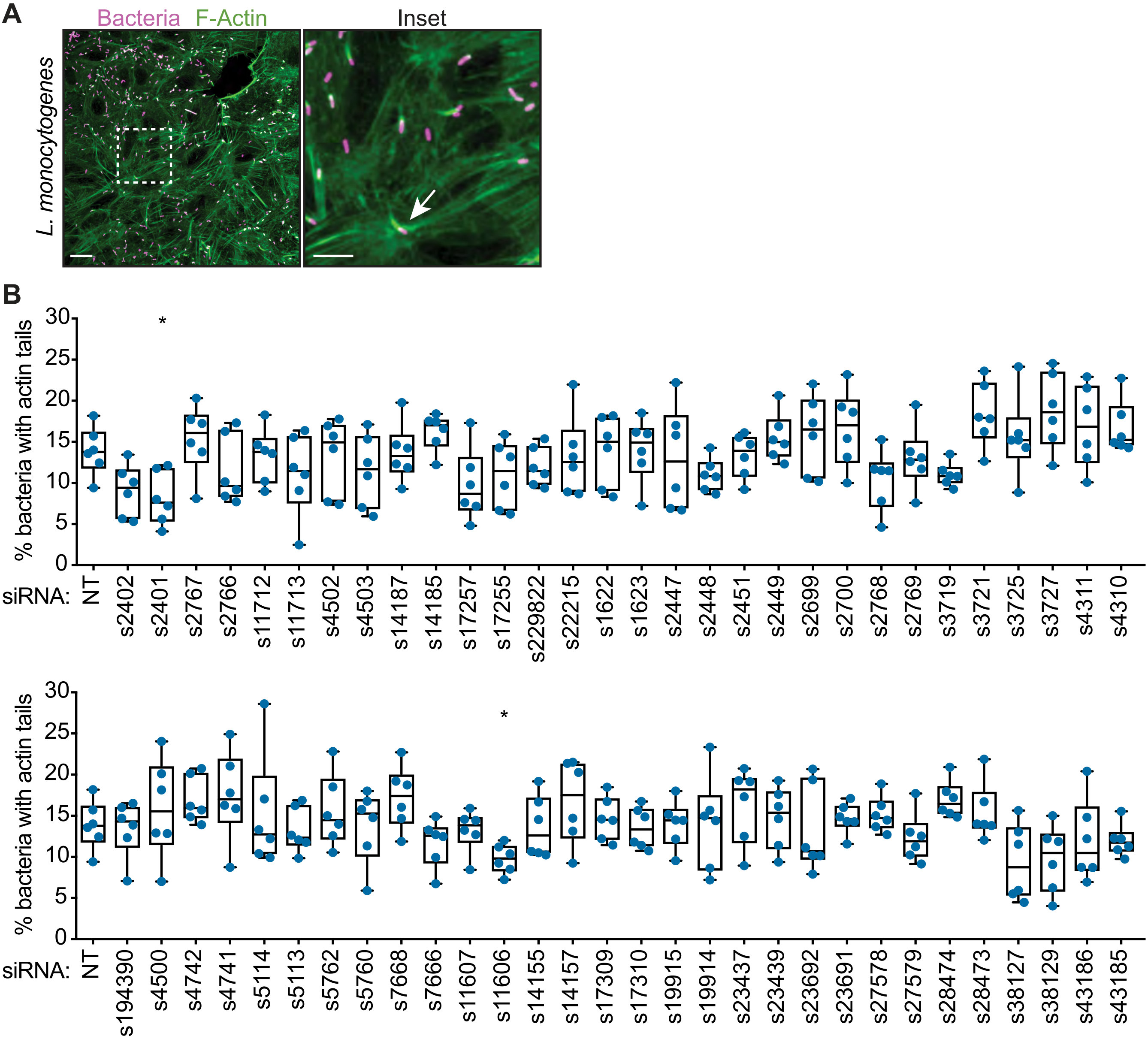
A
Bacteria F-Actin
Inset
L. monocytogenes
B
% bacteria with actin tails
siRNA:
NT s2402 s2401 s2767 s2766 s11712 s11713 s4502 s4503 s14187 s14185 s17257 s17255 s229822 s22215 s1622 s1623 s2447 s2448 s2451 s2449 s2699 s2700 s2768 s2769 s3719 s3721 s3725 s3727 s4311 s4310
NT s194390 s4500 s4742 s4741 s5114 s5113 s5762 s5760 s7668 s7666 s11607 s11606 s14155 s14157 s17309 s17310 s19915 s19914 s23437 s23439 s23692 s23691 s27578 s27579 s28474 s28473 s38127 s38129 s43186 s43185

Fig S1

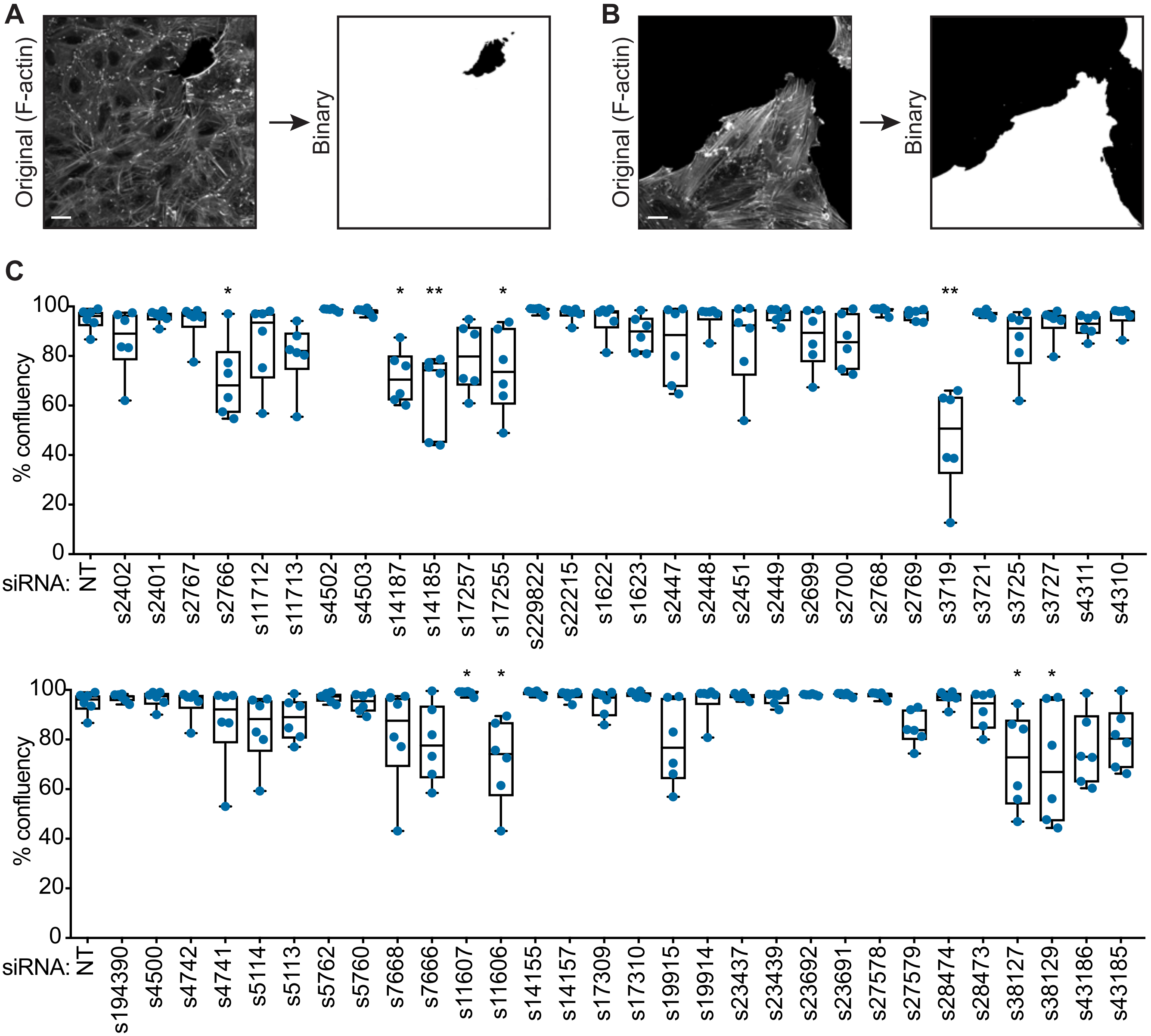

A
Original (F-actin)
Binary
B
Original (F-actin)
Binary
C
% confluency
100
80
60
40
20
0
*
**
*
**
siRNA: NT s2402 s2401 s2767 s2766 s11712 s11713 s4502 s4503 s14187 s14185 s17257 s17255 s229822 s22215 s1622 s1623 s2447 s2448 s2451 s2449 s2699 s2700 s2768 s2769 s3719 s3721 s3725 s3727 s4311 s4310
% confluency
siRNA: NT s194390 s4500 s4742 s4741 s5114 s5113 s5762 s5760 s7668 s7666 s11607 s11606 s14155 s14157 s17309 s17310 s19915 s19914 s23437 s23439 s23692 s23691 s27578 s27579 s28474 s28473 s38127 s38129 s43186 s43185


Fig S2

**Table S1: Primary siRNA screen results**

increased 2+ SD
decreased 2+ SD

| | | | NT Avg: 11.34 *Listeria* A | NT Avg: 12.2 *Listeria* B | NT Avg: 83 *Listeria* A | NT Avg: 85.7 *Listeria* B |
|---|---|---|---|---|---|---|
| **Entrez Gene ID** | **siRNA ID** | **Gene Symbol** | **Avg Cluster Size** | **Avg Cluster Size** | **# Clusters/well** | **# Clusters/well** |
| 10093 | s288550 | ARPC4 | 10.35 | 10.48 | 57 | 65 |
| ArpC4 | s288549 | ARPC4 | 14.73 | 13.74 | 125 | 120 |
| | s288548 | ARPC4 | 8.90 | 9.06 | 67 | 67 |
| 832 | s2403 | CAPZB | 7.66 | 8.33 | 61 | 67 |
| CapzB | s2402 | CAPZB | 9.39 | 8.91 | 94 | 117 |
| | s2401 | CAPZB | 9.55 | 9.51 | 109 | 105 |
| 163 | s38 | AP2B1 | 9.56 | 10.99 | 91 | 86 |
| AP2B1 | s37 | AP2B1 | 9.91 | 11.91 | 91 | 76 |
| | s36 | AP2B1 | 11.37 | 10.42 | 82 | 84 |
| 382 | s1567 | ARF6 | 11.46 | 11.16 | 74 | 79 |
| ARF6 | s1565 | ARF6 | 10.04 | 10.68 | 79 | 68 |
| | s223978 | ARF6 | 10.72 | 11.92 | 72 | 77 |
| 387 | s760 | RHOA | 10.66 | 12.43 | 76 | 83 |
| RhoA | s759 | RHOA | 9.70 | 10.79 | 90 | 80 |
| | s758 | RHOA | 8.36 | 8.80 | 44 | 50 |
| 389 | s99 | RHOC | 8.27 | 9.92 | 55 | 65 |
| RhoC | s98 | RHOC | 10.56 | 9.68 | 88 | 79 |
| | s97 | RHOC | 9.92 | 10.77 | 75 | 74 |
| 408 | s1622 | ARRB1 | 5.65 | 8.83 | 65 | 63 |
| beta-arrestin 1 | s1624 | ARRB1 | 7.35 | 9.45 | 72 | 65 |
| | s1623 | ARRB1 | 6.76 | 6.27 | 42 | 51 |
| 409 | s1627 | ARRB2 | 11.16 | 12.75 | 31 | 48 |
| beta-arrestin2 | s1625 | ARRB2 | 9.39 | 10.19 | 57 | 68 |
| | s1626 | ARRB2 | 11.74 | 11.32 | 77 | 76 |
| 857 | s2447 | CAV1 | 9.18 | 8.30 | 73 | 67 |
| Caveolin 1 | s2446 | CAV1 | 8.64 | 11.83 | 78 | 48 |
| | s2448 | CAV1 | 9.30 | 7.80 | 67 | 97 |
| 858 | s2451 | CAV2 | 7.87 | 8.25 | 103 | 108 |
| Caveolin 2 | s2449 | CAV2 | 8.48 | 9.56 | 91 | 82 |
| | s2450 | CAV2 | 9.71 | 13.45 | 80 | 56 |
| 928 | s2598 | CD9 | 11.84 | 16.02 | 80 | 63 |
| TSPAN-29 | s2597 | CD9 | 12.02 | 11.57 | 89 | 82 |
| | s2599 | CD9 | 10.04 | 9.39 | 69 | 64 |
| 960 | s284408 | CD44 | 8.69 | 10.60 | 55 | 75 |
| CD44 | s307357 | CD44 | 11.80 | 10.16 | 71 | 86 |
| | s284407 | CD44 | 11.27 | 12.66 | 119 | 112 |
| 967 | s2699 | CD63 | 6.97 | 9.53 | 38 | 43 |
| Tspan30/CD63 | s2700 | CD63 | 8.77 | 9.01 | 78 | 81 |
| | s2701 | CD63 | 9.56 | 11.48 | 61 | 69 |
| 975 | s2723 | CD81 | 8.89 | 10.48 | 83 | 82 |
| Tsapn-28/CD81 | s2722 | CD81 | 11.07 | 12.20 | 87 | 80 |
| | s2724 | CD81 | 8.84 | 10.37 | 50 | 59 |
| 977 | s194332 | CD151 | 7.74 | 8.04 | 42 | 57 |
| TSPAN-24 | s194333 | CD151 | 11.79 | 13.24 | 96 | 84 |
| | s2728 | CD151 | 12.04 | 12.67 | 83 | 69 |

increased 2+ SD
decreased 2+ SD

| Entrez Gene ID | siRNA ID | Gene Symbol | Avg Cluster Size (NT Avg: 11.34, *Listeria* A) | Avg Cluster Size (NT Avg: 12.2, *Listeria* B) | # Clusters/well (NT Avg: 83, *Listeria* A) | # Clusters/well (NT Avg: 85.7, *Listeria* B) |
|---|---|---|---|---|---|---|
| 998 | s2767 | CDC42 | 6.97 | 6.69 | 93 | 96 |
| Cdc42 | s2766 | CDC42 | 4.05 | 7.17 | 41 | 52 |
| | s2765 | CDC42 | 8.72 | 10.53 | 93 | 94 |
| 999 | s2768 | CDH1 | 9.50 | 4.67 | 6 | 21 |
| E-cadherin | s2769 | CDH1 | 7.00 | 6.07 | 11 | 42 |
| | s2770 | CDH1 | 9.11 | 7.52 | 35 | 63 |
| 1000 | s2772 | CDH2 | 8.31 | 17.09 | 55 | 67 |
| N-cadherin, type 1 | s2773 | CDH2 | 10.91 | 10.16 | 95 | 112 |
| | s2771 | CDH2 | 8.07 | 10.40 | 102 | 96 |
| 1173 | s3114 | AP2M1 | 8.21 | 9.57 | 107 | 81 |
| AP2-u2 | s3112 | AP2M1 | 9.75 | 11.04 | 84 | 93 |
| | s3113 | AP2M1 | 13.03 | 14.54 | 120 | 98 |
| 1213 | s477 | CLTC | 9.11 | 10.64 | 120 | 95 |
| Clathrin Heavy Chn | s476 | CLTC | 11.67 | 13.67 | 108 | 95 |
| | s475 | CLTC | 13.22 | 14.94 | 114 | 95 |
| 1495 | s3716 | CTNNA1 | 7.23 | 10.65 | 78 | 63 |
| alpha-catenin 1 | s3717 | CTNNA1 | 7.84 | 7.64 | 86 | 114 |
| | s3718 | CTNNA1 | 9.38 | 13.50 | 102 | 74 |
| 1496 | s3719 | CTNNA2 | 6.05 | 6.74 | 38 | 50 |
| alpha-catenin 2 | s3721 | CTNNA2 | 9.48 | 10.24 | 95 | 83 |
| | s3720 | CTNNA2 | 12.34 | 12.80 | 67 | 80 |
| 1499 | s438 | CTNNB1 | 11.09 | 9.18 | 45 | 66 |
| beta-catenin | s437 | CTNNB1 | 11.73 | 10.81 | 60 | 72 |
| | s436 | CTNNB1 | 7.66 | 7.48 | 59 | 63 |
| 1500 | s3726 | CTNND1 | 10.43 | 11.68 | 79 | 72 |
| p120 catenin | s3725 | CTNND1 | 4.95 | 6.50 | 115 | 103 |
| | s3727 | CTNND1 | 6.18 | 8.72 | 40 | 60 |
| 1785 | s285562 | DNM2 | 11.46 | 10.28 | 95 | 96 |
| dynamin-2 | s285561 | DNM2 | 11.86 | 11.16 | 91 | 99 |
| | s285560 | DNM2 | 9.55 | 12.15 | 74 | 89 |
| 1824 | s4311 | DSC2 | 6.27 | 8.39 | 75 | 88 |
| desmocollin-2 | s4309 | DSC2 | 10.81 | 11.55 | 75 | 95 |
| | s4310 | DSC2 | 8.49 | 9.86 | 81 | 97 |
| 1825 | s4312 | DSC3 | 9.41 | 11.23 | 75 | 99 |
| desmocollin-3 | s4313 | DSC3 | 12.39 | 13.46 | 74 | 81 |
| | s4314 | DSC3 | 13.88 | 12.82 | 77 | 76 |
| 1828 | s4321 | DSG1 | 12.26 | 12.75 | 93 | 99 |
| desmoglein-1 | s4323 | DSG1 | 9.92 | 12.68 | 71 | 74 |
| | s4322 | DSG1 | 10.32 | 9.98 | 95 | 90 |
| 1829 | s4326 | DSG2 | 9.56 | 11.30 | 109 | 97 |
| desmoglein-2 | s4324 | DSG2 | 9.91 | 13.15 | 91 | 82 |
| | s4325 | DSG2 | 9.96 | 11.48 | 89 | 92 |
| 1832 | s4335 | DSP | 9.02 | 10.87 | 60 | 63 |
| Desmoplakin | s4334 | DSP | 9.07 | 10.28 | 88 | 87 |
| | s4333 | DSP | 12.85 | 10.41 | 97 | 95 |
| 1943 | s194390 | EFNA2 | 9.13 | 10.20 | 76 | 75 |

| | | | NT Avg: 11.34 *Listeria* A | NT Avg: 12.2 *Listeria* B | NT Avg: 83 *Listeria* A | NT Avg: 85.7 *Listeria* B |
|---|---|---|---|---|---|---|
| increased 2+ SD | | | | | | |
| decreased 2+ SD | | | | | | |
| **Entrez Gene ID** | **siRNA ID** | **Gene Symbol** | **Avg Cluster Size** | **Avg Cluster Size** | **# Clusters/well** | **# Clusters/well** |
| Ephrin-A2 | s223469 | EFNA2 | 8.76 | 11.35 | 114 | 97 |
| | s4500 | EFNA2 | 8.01 | 9.41 | 73 | 92 |
| 1944 | s4501 | EFNA3 | 10.26 | 11.01 | 89 | 107 |
| Ephrin-A3 | s4502 | EFNA3 | 7.68 | 7.48 | 72 | 84 |
| | s4503 | EFNA3 | 8.04 | 7.80 | 69 | 88 |
| 1945 | s371795 | EFNA4 | 10.98 | 13.06 | 101 | 97 |
| Ephrin-A4 | s371794 | EFNA4 | 10.07 | 11.68 | 73 | 76 |
| | s371793 | EFNA4 | 10.84 | 11.02 | 107 | 91 |
| 1946 | s4508 | EFNA5 | 8.46 | 8.49 | 87 | 92 |
| Ephrin-A5 | s4507 | EFNA5 | 11.01 | 11.33 | 93 | 96 |
| | s4509 | EFNA5 | 14.11 | 14.34 | 53 | 50 |
| 1947 | s4511 | EFNB1 | 7.40 | 9.89 | 45 | 63 |
| Ephrin-B1 | s4510 | EFNB1 | 10.68 | 9.63 | 81 | 92 |
| | s4512 | EFNB1 | 11.59 | 11.26 | 74 | 95 |
| 1948 | s4515 | EFNB2 | 8.91 | 9.42 | 96 | 69 |
| Ephrin-B2 | s4513 | EFNB2 | 10.51 | 10.58 | 87 | 101 |
| | s4514 | EFNB2 | 10.22 | 11.83 | 92 | 80 |
| 1969 | s4564 | EPHA2 | 9.18 | 11.57 | 45 | 67 |
| Ephrin type A receptor 2 | s4566 | EPHA2 | 10.62 | 11.98 | 78 | 82 |
| | s4565 | EPHA2 | 12.41 | 9.90 | 74 | 87 |
| 2017 | s345349 | CTTN | 10.37 | 10.26 | 63 | 85 |
| cortactin | s345348 | CTTN | 13.29 | 11.92 | 82 | 95 |
| | s345347 | CTTN | 11.92 | 10.57 | 83 | 89 |
| 2041 | s4721 | EPHA1 | 8.80 | 10.88 | 107 | 80 |
| Ephrin type A receptor 1 | s4719 | EPHA1 | 9.99 | 11.84 | 77 | 91 |
| | s223490 | EPHA1 | 10.37 | 11.73 | 105 | 107 |
| 2042 | s4724 | EPHA3 | 9.17 | 12.47 | 82 | 91 |
| Ephrin type A receptor 3 | s4722 | EPHA3 | 10.76 | 10.24 | 88 | 90 |
| | s4723 | EPHA3 | 8.99 | 10.55 | 91 | 85 |
| 2045 | s4733 | EPHA7 | 8.92 | 11.69 | 48 | 95 |
| Ephrin type A receptor 7 | s4731 | EPHA7 | 8.41 | 10.49 | 76 | 81 |
| | s4732 | EPHA7 | 5.28 | 6.29 | 47 | 68 |
| 2046 | s4736 | EPHA8 | 9.13 | 10.72 | 79 | 92 |
| Ephrin type A receptor 8 | s4735 | EPHA8 | 11.22 | 12.46 | 82 | 100 |
| | s223498 | EPHA8 | 9.38 | 8.53 | 80 | 75 |
| 2048 | s4742 | EPHB2 | 6.97 | 7.76 | 76 | 71 |
| Ephrin type B receptor 2 | s4741 | EPHB2 | 8.80 | 10.31 | 82 | 80 |
| | s4740 | EPHB2 | 12.10 | 13.63 | 98 | 80 |
| 2049 | s4743 | EPHB3 | 11.39 | 11.56 | 70 | 80 |
| Ephrin type B receptor 3 | s4745 | EPHB3 | 11.77 | 12.18 | 82 | 85 |
| | s4744 | EPHB3 | 10.69 | 11.25 | 104 | 102 |
| 2241 | s5110 | FER | 9.67 | 10.14 | 92 | 84 |
| Fer (F-bar) | s5111 | FER | 10.76 | 11.51 | 100 | 90 |
| | s5109 | FER | 10.46 | 10.31 | 48 | 74 |
| 2242 | s5114 | FES | 7.64 | 9.78 | 97 | 77 |
| Fes (F-bar) | s5113 | FES | 8.13 | 9.36 | 100 | 90 |

| | | | NT Avg: 11.34 *Listeria* A | NT Avg: 12.2 *Listeria* B | NT Avg: 83 *Listeria* A | NT Avg: 85.7 *Listeria* B |
|---|---|---|---|---|---|---|
| increased 2+ SD | | | | | | |
| decreased 2+ SD | | | | | | |
| **Entrez Gene ID** | **siRNA ID** | **Gene Symbol** | **Avg Cluster Size** | **Avg Cluster Size** | **# Clusters/well** | **# Clusters/well** |
| | s5112 | FES | 10.31 | 14.08 | 91 | 91 |
| 2319 | s5285 | FLOT2 | 10.92 | 12.39 | 97 | 93 |
| Flotillin 2 | s5286 | FLOT2 | 12.09 | 12.06 | 90 | 96 |
| | s5284 | FLOT2 | 7.95 | 10.04 | 96 | 96 |
| 2697 | s5757 | GJA1 | 9.27 | 12.73 | 98 | 93 |
| Connexin 43 | s5758 | GJA1 | 9.87 | 10.52 | 92 | 95 |
| | s5759 | GJA1 | 11.57 | 13.47 | 84 | 114 |
| 2700 | s5761 | GJA3 | 9.32 | 10.16 | 77 | 89 |
| Connexin 46 | s5762 | GJA3 | 9.52 | 10.33 | 87 | 103 |
| | s5760 | GJA3 | 8.95 | 10.02 | 61 | 83 |
| 2701 | s5763 | GJA4 | 8.71 | 9.36 | 78 | 87 |
| Connexin 37 | s5765 | GJA4 | 8.89 | 10.75 | 96 | 91 |
| | s5764 | GJA4 | 10.88 | 14.08 | 73 | 75 |
| 2702 | s5767 | GJA5 | 10.41 | 9.23 | 85 | 79 |
| Connexin 40 | s5768 | GJA5 | 10.01 | 10.08 | 94 | 87 |
| | s5766 | GJA5 | 11.26 | 11.20 | 90 | 101 |
| 2706 | s5775 | GJB2 | 13.13 | 12.49 | 90 | 102 |
| Connexin 26 | s5777 | GJB2 | 9.75 | 8.85 | 79 | 74 |
| | s5776 | GJB2 | 13.88 | 12.34 | 94 | 116 |
| 2707 | s5779 | GJB3 | 11.03 | 9.68 | 90 | 111 |
| Connexin 31 | s5778 | GJB3 | 12.38 | 9.22 | 90 | 100 |
| | s5780 | GJB3 | 9.20 | 9.17 | 56 | 72 |
| 2709 | s5783 | GJB5 | 11.12 | 10.63 | 106 | 115 |
| Connexin 31.3 | s223727 | GJB5 | 9.77 | 12.46 | 79 | 98 |
| | s5782 | GJB5 | 9.88 | 10.89 | 94 | 101 |
| 3383 | s7087 | ICAM1 | 9.81 | 10.56 | 88 | 71 |
| ICAM1 | s7088 | ICAM1 | 10.20 | 11.83 | 98 | 103 |
| | s7086 | ICAM1 | 7.68 | 8.66 | 37 | 59 |
| 3728 | s7668 | JUP | 8.67 | 9.16 | 95 | 119 |
| gamma-catenin | s7666 | JUP | 7.44 | 9.99 | 59 | 72 |
| | s7667 | JUP | 11.56 | 10.75 | 86 | 113 |
| 4478 | s8985 | MSN | 7.08 | 10.65 | 59 | 74 |
| Moesin | s8984 | MSN | 10.66 | 13.86 | 93 | 95 |
| | s8986 | MSN | 7.78 | 10.03 | 69 | 77 |
| 5110 | s10130 | PCMT1 | 10.23 | 13.83 | 80 | 84 |
| Endophilin-B1 | s10132 | PCMT1 | 10.47 | 12.14 | 108 | 95 |
| | s10131 | PCMT1 | 9.62 | 10.47 | 61 | 85 |
| 5819 | s11607 | PVRL2 | 9.24 | 9.63 | 91 | 95 |
| Nectin 2 | s11606 | PVRL2 | 7.24 | 10.10 | 58 | 60 |
| | s11608 | PVRL2 | 9.99 | 11.26 | 87 | 72 |
| 5868 | s11678 | RAB5A | 11.59 | 11.92 | 80 | 77 |
| Rab5a | s11680 | RAB5A | 11.43 | 12.95 | 69 | 77 |
| | s11679 | RAB5A | 12.89 | 9.28 | 74 | 76 |
| 5879 | s11712 | RAC1 | 6.24 | 7.97 | 45 | 86 |
| Rac1 | s11711 | RAC1 | 4.27 | 3.96 | 52 | 55 |
| | s11713 | RAC1 | 5.15 | 7.38 | 67 | 64 |

| increased 2+ SD / decreased 2+ SD | | | NT Avg: 11.34 *Listeria* A | NT Avg: 12.2 *Listeria* B | NT Avg: 83 *Listeria* A | NT Avg: 85.7 *Listeria* B |
|---|---|---|---|---|---|---|
| **Entrez Gene ID** | **siRNA ID** | **Gene Symbol** | **Avg Cluster Size** | **Avg Cluster Size** | **# Clusters/well** | **# Clusters/well** |
| 5962 | s11899 | RDX | 9.45 | 10.10 | 86 | 102 |
| Radixin | s11900 | RDX | 11.11 | 11.41 | 85 | 98 |
| | s11901 | RDX | 9.77 | 12.42 | 52 | 83 |
| 6455 | s12796 | SH3GL1 | 11.32 | 12.44 | 93 | 91 |
| Endophilin-A2 | s12797 | SH3GL1 | 8.92 | 11.48 | 91 | 75 |
| | s12798 | SH3GL1 | 10.25 | 10.96 | 106 | 100 |
| 7082 | s14155 | TJP1 | 9.33 | 9.86 | 75 | 98 |
| ZO-1 | s14156 | TJP1 | 12.66 | 10.87 | 82 | 99 |
| | s14157 | TJP1 | 9.28 | 10.37 | 82 | 84 |
| 7094 | s14187 | TLN1 | 7.73 | 10.85 | 110 | 122 |
| Talin 1 | s14186 | TLN1 | 5.07 | 8.18 | 91 | 110 |
| | s14185 | TLN1 | 9.58 | 9.55 | 113 | 121 |
| 7106 | s14216 | TSPAN4 | 10.11 | 11.98 | 96 | 80 |
| TSPAN-4/NAG-2 | s14215 | TSPAN4 | 12.30 | 11.51 | 89 | 93 |
| | s14217 | TSPAN4 | 9.74 | 8.86 | 39 | 73 |
| 7251 | s14439 | TSG101 | 11.04 | 14.11 | 94 | 72 |
| Tsg101 (ESCRT-I) | s14440 | TSG101 | 9.27 | 10.77 | 84 | 87 |
| | s14441 | TSG101 | 9.83 | 9.42 | 70 | 88 |
| 7412 | s14759 | VCAM1 | 10.99 | 10.35 | 82 | 68 |
| VCAM-1 | s14761 | VCAM1 | 11.83 | 12.65 | 95 | 97 |
| | s14760 | VCAM1 | 10.26 | 11.03 | 98 | 93 |
| 7414 | s14762 | VCL | 9.12 | 10.14 | 85 | 91 |
| Vinculin | s14763 | VCL | 10.83 | 11.79 | 88 | 103 |
| | s14764 | VCL | 11.55 | 12.37 | 91 | 90 |
| 7430 | s14796 | EZR | 11.38 | 12.09 | 98 | 96 |
| Ezrin | s14795 | EZR | 10.92 | 10.81 | 95 | 101 |
| | s14797 | EZR | 9.96 | 11.91 | 104 | 94 |
| 8436 | s16008 | SDPR | 10.30 | 9.40 | 89 | 105 |
| Cavin-2 | s16006 | SDPR | 10.06 | 9.52 | 66 | 94 |
| | s16007 | SDPR | 10.90 | 10.67 | 94 | 93 |
| 8522 | s16201 | GAS7 | 9.15 | 8.69 | 81 | 85 |
| GSA7 (F-bar) | s16202 | GAS7 | 9.27 | 11.48 | 74 | 63 |
| | s16200 | GAS7 | 10.18 | 9.20 | 89 | 90 |
| 8976 | s17134 | WASL | 11.42 | 12.10 | 113 | 111 |
| N-WASP | s17133 | WASL | 8.35 | 10.94 | 113 | 95 |
| | s17132 | WASL | 7.07 | 11.39 | 75 | 74 |
| 9050 | s17253 | PSTPIP2 | 10.07 | 10.75 | 71 | 72 |
| PSTPIP2 | s17254 | PSTPIP2 | 12.99 | 14.48 | 73 | 97 |
| | s17252 | PSTPIP2 | 11.68 | 12.78 | 104 | 91 |
| 9051 | s17257 | PSTPIP1 | 8.64 | 8.04 | 69 | 74 |
| PSTPIP1 (F-bar) | s17255 | PSTPIP1 | 6.20 | 7.64 | 45 | 53 |
| | s17256 | PSTPIP1 | 9.87 | 10.79 | 124 | 99 |
| 9069 | s17298 | CLDN12 | 9.02 | 10.52 | 83 | 89 |
| Claudin-12 | s17297 | CLDN12 | 11.04 | 10.81 | 91 | 75 |
| | s17299 | CLDN12 | 10.97 | 9.87 | 64 | 90 |
| 9074 | s17309 | CLDN6 | 8.66 | 9.41 | 79 | 83 |

| | | | NT Avg: 11.34 *Listeria* A | NT Avg: 12.2 *Listeria* B | NT Avg: 83 *Listeria* A | NT Avg: 85.7 *Listeria* B |
|---|---|---|---|---|---|---|
| increased 2+ SD | | | | | | |
| decreased 2+ SD | | | | | | |
| **Entrez Gene ID** | **siRNA ID** | **Gene Symbol** | **Avg Cluster Size** | **Avg Cluster Size** | **# Clusters/well** | **# Clusters/well** |
| Claudin-6 | s17311 | CLDN6 | 11.15 | 13.48 | 87 | 84 |
| | s17310 | CLDN6 | 10.07 | 8.93 | 96 | 98 |
| 9075 | s17314 | CLDN2 | 7.72 | 11.25 | 106 | 107 |
| Claudin-2 | s17312 | CLDN2 | 8.31 | 10.45 | 83 | 99 |
| | s225076 | CLDN2 | 8.66 | 9.21 | 71 | 73 |
| 9076 | s17316 | CLDN1 | 10.50 | 11.92 | 86 | 98 |
| Claudin-1 | s17317 | CLDN1 | 10.49 | 10.66 | 103 | 97 |
| | s17315 | CLDN1 | 8.49 | 11.04 | 94 | 113 |
| 9080 | s194981 | CLDN9 | 10.78 | 11.73 | 92 | 92 |
| Claudin-9 | s194980 | CLDN9 | 9.90 | 13.36 | 78 | 66 |
| | s194979 | CLDN9 | 9.03 | 10.94 | 73 | 78 |
| 9146 | s17480 | HGS | 9.73 | 10.37 | 71 | 83 |
| Hrs | s17481 | HGS | 9.68 | 11.43 | 59 | 92 |
| | s17482 | HGS | 10.26 | 11.08 | 87 | 83 |
| 9322 | s17816 | TRIP10 | 9.25 | 12.31 | 91 | 86 |
| CIP4 | s17815 | TRIP10 | 11.12 | 10.57 | 85 | 93 |
| | s17814 | TRIP10 | 12.78 | 11.09 | 94 | 99 |
| 9525 | s18274 | VPS4B | 10.91 | 14.45 | 91 | 98 |
| SKD1 | s18273 | VPS4B | 8.02 | 9.85 | 109 | 102 |
| | s18272 | VPS4B | 12.56 | 11.38 | 100 | 104 |
| 9788 | s18916 | MTSS1 | 9.71 | 12.31 | 87 | 91 |
| MIM | s18917 | MTSS1 | 11.14 | 11.06 | 88 | 107 |
| | s18915 | MTSS1 | 8.79 | 10.08 | 73 | 79 |
| 10052 | s19541 | GJC1 | 10.43 | 9.82 | 101 | 104 |
| Connexin-45 | s19543 | GJC1 | 9.69 | 11.10 | 74 | 88 |
| | s223072 | GJC1 | 5.03 | 8.48 | 79 | 85 |
| 10211 | s19915 | FLOT1 | 6.71 | 9.48 | 78 | 79 |
| Flotillin 1 | s19914 | FLOT1 | 8.67 | 7.75 | 51 | 97 |
| | s19913 | FLOT1 | 8.57 | 10.48 | 72 | 93 |
| 10458 | s20464 | BAIAP2 | 8.23 | 10.77 | 92 | 93 |
| IRSp53 (I-bar) | s20465 | BAIAP2 | 10.00 | 13.23 | 73 | 75 |
| | s20463 | BAIAP2 | 9.93 | 8.66 | 73 | 102 |
| 10804 | s21231 | GJB6 | 9.06 | 10.12 | 100 | 86 |
| Connexin 30 | s21232 | GJB6 | 11.37 | 12.49 | 82 | 98 |
| | s21233 | GJB6 | 11.62 | 11.86 | 74 | 81 |
| 11252 | s229822 | PACSIN2 | 10.11 | 9.39 | 90 | 76 |
| Pacsin2 (F-bar) | s22216 | PACSIN2 | 11.15 | 9.92 | 88 | 88 |
| | s22215 | PACSIN2 | 6.23 | 9.89 | 43 | 64 |
| 11267 | s22248 | SNF8 | 9.79 | 10.91 | 107 | 92 |
| EAP30 (Escrt-II) | s22247 | SNF8 | 10.05 | 9.32 | 96 | 92 |
| | s22249 | SNF8 | 12.51 | 14.33 | 110 | 99 |
| 23048 | s22914 | FNBP1 | 10.24 | 11.93 | 59 | 75 |
| FBP17 | s22915 | FNBP1 | 9.94 | 10.62 | 90 | 84 |
| | s22916 | FNBP1 | 10.58 | 10.46 | 96 | 115 |
| 23268 | s23437 | DNMBP | 9.71 | 9.59 | 89 | 74 |
| Tuba (Bar domain) | s23438 | DNMBP | 11.50 | 11.83 | 102 | 106 |

| increased 2+ SD | | | NT Avg: 11.34 *Listeria* A | NT Avg: 12.2 *Listeria* B | NT Avg: 83 *Listeria* A | NT Avg: 85.7 *Listeria* B |
|---|---|---|---|---|---|---|
| decreased 2+ SD | | | | | | |
| **Entrez Gene ID** | **siRNA ID** | **Gene Symbol** | **Avg Cluster Size** | **Avg Cluster Size** | **# Clusters/well** | **# Clusters/well** |
| | s23439 | DNMBP | 9.18 | 9.33 | 92 | 76 |
| 23380 | s23693 | SRGAP2 | 11.51 | 12.50 | 89 | 80 |
| srGAP2 (F-bar) | s23692 | SRGAP2 | 9.67 | 9.99 | 82 | 98 |
| | s23691 | SRGAP2 | 9.57 | 10.38 | 88 | 92 |
| 23607 | s24191 | CD2AP | 9.56 | 12.64 | 89 | 104 |
| CD2AP | s24192 | CD2AP | 8.26 | 13.13 | 76 | 72 |
| | s24193 | CD2AP | 7.59 | 11.82 | 68 | 72 |
| 25945 | s24804 | PVRL3 | 9.91 | 12.86 | 76 | 86 |
| Nectin-3 | s24803 | PVRL3 | 11.18 | 13.37 | 112 | 108 |
| | s24802 | PVRL3 | 11.22 | 12.19 | 91 | 89 |
| 26052 | s25018 | DNM3 | 10.08 | 11.78 | 96 | 81 |
| Dynamin-3 | s25017 | DNM3 | 13.94 | 12.97 | 95 | 94 |
| | s25019 | DNM3 | 8.15 | 13.25 | 47 | 60 |
| 29763 | s26562 | PACSIN3 | 11.02 | 10.73 | 81 | 86 |
| Pacsin3 (F-bar) | s26563 | PACSIN3 | 10.21 | 10.44 | 81 | 103 |
| | s26564 | PACSIN3 | 6.88 | 8.06 | 25 | 50 |
| 50807 | s27099 | ASAP1 | 14.04 | 13.65 | 85 | 106 |
| ASAP1 | s27101 | ASAP1 | 12.25 | 14.79 | 79 | 84 |
| | s27100 | ASAP1 | 8.03 | 9.24 | 77 | 80 |
| 50848 | s27150 | F11R | 9.92 | 13.40 | 72 | 77 |
| JAM-A | s27151 | F11R | 10.09 | 11.03 | 69 | 86 |
| | s27152 | F11R | 8.39 | 8.87 | 57 | 89 |
| 51160 | s27577 | VPS28 | 10.86 | 10.68 | 91 | 107 |
| hVps28 (Escrt-1) | s27578 | VPS28 | 9.09 | 10.19 | 82 | 93 |
| | s27579 | VPS28 | 8.44 | 8.87 | 75 | 91 |
| 51294 | s27870 | PCDH12 | 9.99 | 14.64 | 96 | 86 |
| VE-Cadherin | s27869 | PCDH12 | 11.29 | 12.51 | 94 | 93 |
| | s27871 | PCDH12 | 8.74 | 12.72 | 91 | 98 |
| 51652 | s28475 | VPS24 | 10.13 | 11.08 | 90 | 99 |
| CHMP3 (ESCRT-III) | s28474 | VPS24 | 14.53 | 15.25 | 96 | 105 |
| | s28473 | VPS24 | 13.35 | 15.48 | 86 | 94 |
| 54874 | s29645 | FNBP1L | 10.73 | 12.82 | 86 | 105 |
| TOCA1 | s29646 | FNBP1L | 10.82 | 10.75 | 96 | 92 |
| | s29644 | FNBP1L | 8.88 | 10.91 | 86 | 78 |
| 56904 | s32358 | SH3GLB2 | 10.82 | 11.27 | 90 | 75 |
| Endophilin-B2 | s32357 | SH3GLB2 | 8.69 | 10.80 | 84 | 74 |
| | s32359 | SH3GLB2 | 10.50 | 12.01 | 111 | 103 |
| 57522 | s33218 | SRGAP1 | 4.63 | 5.24 | 73 | 76 |
| srGAP1 | s33219 | SRGAP1 | 13.35 | 11.33 | 99 | 94 |
| | s33217 | SRGAP1 | 13.89 | 10.29 | 94 | 102 |
| 58494 | s33856 | JAM2 | 11.82 | 18.52 | 89 | 81 |
| JAM-B | s33857 | JAM2 | 9.88 | 12.90 | 91 | 88 |
| | s33855 | JAM2 | 11.52 | 12.82 | 95 | 85 |
| 83660 | s38090 | TLN2 | 9.95 | 9.82 | 85 | 98 |
| Talin-2 | s38088 | TLN2 | 8.99 | 11.16 | 69 | 68 |
| | s38089 | TLN2 | 9.12 | 10.65 | 94 | 97 |

| increased 2+ SD | | | NT Avg: 11.34 *Listeria* A | NT Avg: 12.2 *Listeria* B | NT Avg: 83 *Listeria* A | NT Avg: 85.7 *Listeria* B |
|---|---|---|---|---|---|---|
| decreased 2+ SD | | | | | | |
| **Entrez Gene ID** | **siRNA ID** | **Gene Symbol** | **Avg Cluster Size** | **Avg Cluster Size** | **# Clusters/well** | **# Clusters/well** |
| 83700 | s38127 | JAM3 | 7.90 | 10.36 | 48 | 45 |
| JAM-C | s38129 | JAM3 | 6.68 | 8.27 | 37 | 55 |
| | s38128 | JAM3 | 12.20 | 13.11 | 105 | 93 |
| 92154 | s40868 | MTSS1L | 10.18 | 10.52 | 57 | 69 |
| ABBA | s40867 | MTSS1L | 10.70 | 11.16 | 98 | 92 |
| | s40869 | MTSS1L | 9.24 | 8.46 | 66 | 56 |
| 115677 | s41845 | NOSTRIN | 10.88 | 10.96 | 75 | 75 |
| Nostrin | s41847 | NOSTRIN | 11.05 | 12.64 | 111 | 80 |
| | s41846 | NOSTRIN | 10.91 | 13.49 | 87 | 99 |
| 125111 | s42859 | GJD3 | 10.94 | 10.27 | 70 | 70 |
| Connexin 31.9 | s42857 | GJD3 | 10.36 | 10.21 | 78 | 80 |
| | s42858 | GJD3 | 11.19 | 9.25 | 59 | 92 |
| 127534 | s43186 | GJB4 | 8.89 | 6.38 | 55 | 52 |
| Connexin 30.3 | s43185 | GJB4 | 7.39 | 10.41 | 62 | 76 |
| | s43184 | GJB4 | 10.00 | 10.99 | 77 | 99 |
| 147409 | s44997 | DSG4 | 10.79 | 11.67 | 81 | 99 |
| desmoglein-4 | s44999 | DSG4 | 11.78 | 11.85 | 54 | 78 |
| | s44998 | DSG4 | 11.25 | 10.94 | 81 | 97 |
| 284119 | s49506 | PTRF | 14.22 | 14.20 | 96 | 109 |
| Cavin-1 | s49507 | PTRF | 7.94 | 9.32 | 31 | 68 |
| | s49508 | PTRF | 10.05 | 10.82 | 55 | 68 |
| 100506658 | s446558 | OCLN | 12.00 | 11.89 | 107 | 94 |
| Occludin | s446559 | OCLN | 14.04 | 13.94 | 76 | 98 |
| | s446560 | OCLN | 11.08 | 13.78 | 106 | 97 |

Table S2

| RefSeq Accession Number | Gene Symbol | Full Gene Name | Gene ID | siRNA ID | Sense siRNA Sequence | Antisense siRNA Sequence |
|---|---|---|---|---|---|---|
| AK074205 | ARPC4 | actin related protein 2/3 complex, subunit 4, | 10093 | n288550 | UCAACAAUCAUUUACUUAUtt | AUAAGUAAAUGAUUGUUGAgt |
| AK074205 | ARPC4 | actin related protein 2/3 complex, subunit 4, | 10093 | n288549 | AAAUUCAGGUGCAUCCAAAtt | UUUGGAUGCACCUGAAUUUgt |
| AK074205 | ARPC4 | actin related protein 2/3 complex, subunit 4, | 10093 | n288548 | GGUCUAAGAAAGUACCAUAtt | UAUGGUACUUUCUUAGACCtc |
| NM_004930 | CAPZB | capping protein (actin filament) muscle Z-line, | 832 | s2403 | GAAGUACGCUGAACGAGAUtt | AUCUCGUUCAGCGUACUUCtg |
| NM_004930 | CAPZB | capping protein (actin filament) muscle Z-line, | 832 | s2402 | GACUUUUGCAGACAAAUCAtt | UGAUUUGUCUGCAAAAGUCtg |
| NM_004930 | CAPZB | capping protein (actin filament) muscle Z-line, | 832 | s2401 | CGAGAUCUACUUUGGAAAAtt | UUUUCCAAAGUAGAUCUCGtt |
| NM_001030006 | AP2B1 | adaptor-related protein complex 2, beta 1 | 163 | s38 | GGCUUGGAGAUUUCCGGAAtt | UUCCGGAAAUCUCCAAGCCtt |
| NM_001030006 | AP2B1 | adaptor-related protein complex 2, beta 1 | 163 | s37 | GGAGGGUUUUCACGAUGAAtt | UUCAUCGUGAAAACCCUCCag |
| NM_001030006 | AP2B1 | adaptor-related protein complex 2, beta 1 | 163 | s36 | GGAUCCCUAUGUUCGGAAAtt | UUUCCGAACAUAGGGAUCCtc |
| NM_001663 | ARF6 | ADP-ribosylation factor 6 | 382 | s1567 | CCAAGGUCUCAUCUUCGUAtt | UACGAAGAUGAGACCUUGGgt |
| NM_001663 | ARF6 | ADP-ribosylation factor 6 | 382 | s1565 | GUCUCAUCUUCGUAGUGGAtt | UCCACUACGAAGAUGAGACct |
| NM_001663 | ARF6 | ADP-ribosylation factor 6 | 382 | s223978 | CUAAGACCAGUAUAGUAAAtt | UUUACUAUACUGGUCUUAGat |
| NM_001664 | RHOA | ras homolog family member A | 387 | s760 | GGCUUUACUCCGUAACAGAtt | UCUGUUACGGAGUAAAGCCct |
| NM_001664 | RHOA | ras homolog family member A | 387 | s759 | CUAUGAUUAUUAACGAUGUtt | ACAUCGUUAAUAAUCAUAGtt |
| NM_001664 | RHOA | ras homolog family member A | 387 | s758 | CACAGUGUUUGAGAACUAUtt | AUAGUUCUCAAACACUGUGgg |
| NM_001042678 | RHOC | ras homolog family member C | 389 | s99 | AGGACUGCAUUGUUUUCUAtt | UAGAAAACAAUGCAGUCCUgg |
| NM_001042678 | RHOC | ras homolog family member C | 389 | s98 | CACCAGCACUUUAUACACUtt | AGUGUAUAAAGUGCUGGUGtg |
| NM_001042678 | RHOC | ras homolog family member C | 389 | s97 | GCAAGGAUCAGUUUCCGGAtt | UCCGGAAACUGAUCCUUGCtg |
| NM_004041 | ARRB1 | arrestin, beta 1 | 408 | s1622 | GGCUUGCGGUGUGGACUAUtt | AUAGUCCACACCGCAAGCCtt |
| NM_004041 | ARRB1 | arrestin, beta 1 | 408 | s1624 | CCAAUCUCAUAGAACUUGAtt | UCAAGUUCUAUGAGAUUGGta |
| NM_004041 | ARRB1 | arrestin, beta 1 | 408 | s1623 | GGAGAUCUAUUACCAUGGAtt | UCCAUGGUAAUAGAUCUCCtt |
| NM_004313 | ARRB2 | arrestin, beta 2 | 409 | s1627 | AGCCUUCUGUGCUAAAUCAtt | UGAUUUAGCACAGAAGGCUcg |
| NM_004313 | ARRB2 | arrestin, beta 2 | 409 | s1625 | AAGUCUCUGUGAGACAGUAtt | UACUGUCUCACAGAGACUUtg |
| NM_004313 | ARRB2 | arrestin, beta 2 | 409 | s1626 | CGAACAAGAUGACCAGGUAtt | UACCUGGUCAUCUUGUUCGag |
| NM_001753 | CAV1 | caveolin 1, caveolae protein, 22kDa | 857 | s2447 | GCCGUGUCUAUUCCAUCUAtt | UAGAUGGAAUAGACACGGCtg |
| NM_001753 | CAV1 | caveolin 1, caveolae protein, 22kDa | 857 | s2446 | GCUUCCUGAUUGAGAUUCAtt | UGAAUCUCAAUCAGGAAGCtc |
| NM_001753 | CAV1 | caveolin 1, caveolae protein, 22kDa | 857 | s2448 | CCUUCACUGUGACGAAAUAtt | UAUUUCGUCACAGUGAAGGtg |
| NM_001233 | CAV2 | caveolin 2 | 858 | s2451 | AGACCUGCCUAAUGGUUCUtt | AGAACCAUUAGGCAGGUCUtt |
| NM_001233 | CAV2 | caveolin 2 | 858 | s2449 | AUGUUAUCAUUGCUCCAUUtt | AAUGGAGCAAUGAUAACAUct |
| NM_001233 | CAV2 | caveolin 2 | 858 | s2450 | CGGCUCAACUCGCAUCUCAtt | UGAGAUGCGAGUUGAGCCGgt |
| NM_001769 | CD9 | CD9 molecule | 928 | s2598 | UCUUCGAGCAAGAAACUAAtt | UUAGUUUCUUGCUCGAAGAtg |
| NM_001769 | CD9 | CD9 molecule | 928 | s2597 | AGAGGUCUUCGACAAUAAAtt | UUUAUUGUCGAAGACCUCUtt |
| NM_001769 | CD9 | CD9 molecule | 928 | s2599 | CAGUUUAUCUCAGACAUCUtt | AGAUGUCUGAGAUAAACUGtt |
| AF086543 | CD44 | CD44 molecule (Indian blood group) | 960 | n284408 | UGAAGCAUUUAGAAUCAUAtt | UAUGAUUCUAAAUGCUUCAgt |
| AK123567 | CD44 | CD44 molecule (Indian blood group) | 960 | n307357 | GAUUCUCUUGAGUAGUAAAtt | UUUACUACUCAAGAGAAUCat |
| AF086543 | CD44 | CD44 molecule (Indian blood group) | 960 | n284407 | UACACUCCUUAGUUCUUUAtt | UAAAGAACUAAGGAGUGUAag |
| NM_001040034 | CD63 | CD63 molecule | 967 | s2699 | AGAAUUACCCGAAAAACAAtt | UUGUUUUUCGGGUAAUUCUcc |
| NM_001040034 | CD63 | CD63 molecule | 967 | s2700 | GGAUGCAGGCAGAUUUUAAtt | UUAAAAUCUGCCUGCAUCCtg |
| NM_001040034 | CD63 | CD63 molecule | 967 | s2701 | GGAUUAAUUUCAACGAGAAtt | UUCUCGUUGAAAUUAAUCCca |
| NM_004356 | CD81 | CD81 molecule | 975 | s2723 | GAACUUUCCUGUUACCUUUtt | AAAGGUAACAGGAAAGUUCag |
| NM_004356 | CD81 | CD81 molecule | 975 | s2722 | GCCCAACACCUUCUAUGUAtt | UACAUAGAAGGUGUUGGGCgc |
| NM_004356 | CD81 | CD81 molecule | 975 | s2724 | CCACCUCAGUGCUCAAGAAtt | UUCUUGAGCACUGAGGUGGtc |
| NM_001039490 | CD151 | CD151 molecule (Raph blood group) | 977 | s194332 | CCCAACUACUGAGCUGAGAtt | UCUCAGCUCAGUAGUUGGGtg |
| NM_001039490 | CD151 | CD151 molecule (Raph blood group) | 977 | s194333 | GCCUUUUGCUGCGCACCAAtt | UUGGUGCGCAGCAAAAGGCac |
| NM_001039490 | CD151 | CD151 molecule (Raph blood group) | 977 | s2728 | CUGCUGCGCCUGUACUUCAtt | UGAAGUACAGGCGCAGCAGgt |
| NM_001039802 | CDC42 | cell division cycle 42 (GTP binding protein, | 998 | s2767 | CAGUUAUGAUUGGUGGAGAtt | UCUCCACCAAUCAUAACUGtg |
| NM_001039802 | CDC42 | cell division cycle 42 (GTP binding protein, | 998 | s2766 | UGAGAUAACUCACCACUGUtt | ACAGUGGUGAGUUAUCUCAgg |
| NM_001039802 | CDC42 | cell division cycle 42 (GTP binding protein, | 998 | s2765 | UGGUGCUGUUGGUAAAACAtt | UGUUUUACCAACAGCACCAtc |
| NM_004360 | CDH1 | cadherin 1, type 1, E-cadherin (epithelial) | 999 | s2768 | GAACUAUAUUCUUCUGUGAtt | UCACAGAAGAAUAUAGUUCga |
| NM_004360 | CDH1 | cadherin 1, type 1, E-cadherin (epithelial) | 999 | s2769 | CAACAGCUGUGAUCACAGUtt | ACUGUGAUCACAGCUGUUGct |
| NM_004360 | CDH1 | cadherin 1, type 1, E-cadherin (epithelial) | 999 | s2770 | CGUAUACCCUGGUGGUUCAtt | UGAACCACCAGGGUAUACGta |
| NM_001792 | CDH2 | cadherin 2, type 1, N-cadherin (neuronal) | 1000 | s2772 | GGGUAAUCCUCCCAAAUCAtt | UGAUUUGGGAGGAUUACCCga |
| NM_001792 | CDH2 | cadherin 2, type 1, N-cadherin (neuronal) | 1000 | s2773 | GAACAUAUGUGAUGACCGUtt | ACGGUCAUCACAUAUGUUCca |
| NM_001792 | CDH2 | cadherin 2, type 1, N-cadherin (neuronal) | 1000 | s2771 | GUGCAACAGUAUACGUUAAtt | UUAACGUAUACUGUUGCACtt |
| NM_001025205 | AP2M1 | adaptor-related protein complex 2, mu 1 | 1173 | s3114 | GAAGAGCAGUCACAGAUCAtt | UGAUCUGUGACUGCUCUUCtt |
| NM_001025205 | AP2M1 | adaptor-related protein complex 2, mu 1 | 1173 | s3112 | GGAUGAGAUUCUAGACUUUtt | AAAGUCUAGAAUCUCAUCCag |
| NM_001025205 | AP2M1 | adaptor-related protein complex 2, mu 1 | 1173 | s3113 | GGCGAGAGGGUAUCAAGUAtt | UACUUGAUACCCUCUCGCCgc |
| NM_004859 | CLTC | clathrin, heavy chain (Hc) | 1213 | s477 | GGGAAUUCUUCGUACUCCAtt | UGGAGUACGAAGAAUUCCCtt |
| NM_004859 | CLTC | clathrin, heavy chain (Hc) | 1213 | s476 | CGGUUGCUCUUGUUACGGAtt | UCCGUAACAAGAGCAACCGta |
| NM_004859 | CLTC | clathrin, heavy chain (Hc) | 1213 | s475 | GGUUGCUCUUGUUACGGAUtt | AUCCGUAACAAGAGCAACCgt |
| NM_001903 | CTNNA1 | catenin (cadherin-associated protein), alpha 1, | 1495 | s3716 | GUGGAUAAGCUGAACAUUAtt | UAAUGUUCAGCUUAUCCACtt |
| NM_001903 | CTNNA1 | catenin (cadherin-associated protein), alpha 1, | 1495 | s3717 | GUGUAAAGCUUGUUCGAAUtt | AUUCGAACAAGCUUUACACct |
| NM_001903 | CTNNA1 | catenin (cadherin-associated protein), alpha 1, | 1495 | s3718 | CUACGUCGCCUCUACCAAAtt | UUUGGUAGAGGCGACGUAGga |
| NM_004389 | CTNNA2 | catenin (cadherin-associated protein), alpha 2 | 1496 | s3719 | CAAUGUUCCUUUGCUAGUUtt | AACUAGCAAAGGAACAUUGgt |
| NM_004389 | CTNNA2 | catenin (cadherin-associated protein), alpha 2 | 1496 | s3721 | CAGCGGCAAUGAUAUCAUUtt | AAUGAUAUCAUUGCCGCUGtc |
| NM_004389 | CTNNA2 | catenin (cadherin-associated protein), alpha 2 | 1496 | s3720 | CGCUUAUCCAGGCAGCUAAtt | UUAGCUGCCUGGAUAAGCGat |
| NM_001904 | CTNNB1 | catenin (cadherin-associated protein), beta 1, | 1499 | s438 | CUGUUGGAUUGAUUCGAAAtt | UUUCGAAUCAAUCCAACAGta |
| NM_001904 | CTNNB1 | catenin (cadherin-associated protein), beta 1, | 1499 | s437 | GGAUGUUCACAACCGAAUUtt | AAUUCGGUUGUGAACAUCCcg |
| NM_001904 | CTNNB1 | catenin (cadherin-associated protein), beta 1, | 1499 | s436 | GGACCUAUACUUACGAAAAtt | UUUUCGUAAGUAUAGGUCCtc |
| NM_001085458 | CTNND1 | catenin (cadherin-associated protein), delta 1 | 1500 | s3726 | CAUUGACCGGAACCAAAAAtt | UUUUUGGUUCCGGUCAAUGag |
| NM_001085458 | CTNND1 | catenin (cadherin-associated protein), delta 1 | 1500 | s3725 | GCUCGCAACAAAGAAUUAAtt | UUAAUUCUUUGUUGCGAGCat |
| NM_001085458 | CTNND1 | catenin (cadherin-associated protein), delta 1 | 1500 | s3727 | GAACUUAUCAUAUCAAGUUtt | AACUUGAUAUGAUAAGUUCcg |
| AK092344 | DNM2 | dynamin 2 | 1785 | n285562 | ACUGUUUGUUGAAACCAGAtt | UCUGGUUUCAACAAACAGUtg |
| AK092344 | DNM2 | dynamin 2 | 1785 | n285561 | GCACUGGGCCGUCCUUAUAtt | UAUAAGGACGGCCCAGUGCtt |
| AK092344 | DNM2 | dynamin 2 | 1785 | n285560 | GGAGGGAGAGUUUCAGCAAtt | UUGCUGAAACUCUCCCUCCct |
| NM_024422 | DSC2 | desmocollin 2 | 1824 | s4311 | GGUUCAGUCUAUACAACAAtt | UUGUUGUAUAGACUGAACCat |
| NM_024422 | DSC2 | desmocollin 2 | 1824 | s4309 | GACUAUGUCCUGACAUAUAtt | UAUAUGUCAGGACAUAGUCtt |
| NM_024422 | DSC2 | desmocollin 2 | 1824 | s4310 | GCUCAUAUGUAGUACCUAUtt | AUAGGUACUACAUAUGAGCca |
| NM_001941 | DSC3 | desmocollin 3 | 1825 | s4312 | GACGUGGAGUUGAUAAAGAtt | UCUUUAUCAACUCCACGUCca |
| NM_001941 | DSC3 | desmocollin 3 | 1825 | s4313 | CCCAAUAUGUGCGGAUUAAtt | UUAAUCCGCACAUAUUGGGct |

| | | | | | | |
|---|---|---|---|---|---|---|
| NM_001941 | DSC3 | desmocollin 3 | 1825 | s4314 | GAUCAACGGCUAUAAGGCAtt | UGCCUUAUAGCCGUUGAUCtt |
| NM_001942 | DSG1 | desmoglein 1 | 1828 | s4321 | GAACGAUUCUCUCUAUAGAtt | UCUAUAGAGAGAAUCGUUCct |
| NM_001942 | DSG1 | desmoglein 1 | 1828 | s4323 | CAUCCAUAGUUGAUCGAGAtt | UCUCGAUCAACUAUGGAUGtt |
| NM_001942 | DSG1 | desmoglein 1 | 1828 | s4322 | CUGACGCAGAUGAACCGAAtt | UUCGGUUCAUCUGCGUCAGta |
| NM_001943 | DSG2 | desmoglein 2 | 1829 | s4326 | CCAGCAUUCUUGAUCGAGAtt | UCUCGAUCAAGAAUGCUGGta |
| NM_001943 | DSG2 | desmoglein 2 | 1829 | s4324 | CUGUUGCACUGAACGAAGAtt | UCUUCGUUCAGUGCAACAGca |
| NM_001943 | DSG2 | desmoglein 2 | 1829 | s4325 | CAAUACCCUGAAUUCGAAAtt | UUUCGAAUUCAGGGUAUUGgg |
| NM_001008844 | DSP | desmoplakin | 1832 | s4335 | CCGACUGACUUAUGAGAUUtt | AAUCUCAUAAGUCAGUCGGgt |
| NM_001008844 | DSP | desmoplakin | 1832 | s4334 | GGACCUCAUUGACUUCGAUtt | AUCGAAGUCAAUGAGGUCCcg |
| NM_001008844 | DSP | desmoplakin | 1832 | s4333 | GGAACGGUGCAGAAUCGAAtt | UUCGAUUCUGCACCGUUCCtt |
| NM_001405 | EFNA2 | ephrin-A2 | 1943 | s194390 | CCACGAGUAUUACUACAUCtt | GAUGUAGUAAUACUCGUGGcc |
| NM_001405 | EFNA2 | ephrin-A2 | 1943 | s223469 | GGAGCACUACGUGCUGUACtt | GUACAGCACGUAGUGCUCCat |
| NM_001405 | EFNA2 | ephrin-A2 | 1943 | s4500 | CGAGUAUUACUACAUCUCUtt | AGAGAUGUAGUAAUACUCGtg |
| NM_004952 | EFNA3 | ephrin-A3 | 1944 | s4501 | GGAUUGUGGUUUGGAUUGAtt | UCAAUCCAAACCACAAUCCtg |
| NM_004952 | EFNA3 | ephrin-A3 | 1944 | s4502 | ACACUUUUGGAGACCGUAAtt | UUACGGUCUCCAAAAGUGUct |
| NM_004952 | EFNA3 | ephrin-A3 | 1944 | s4503 | AGGUGAACGUGAACGACUAtt | UAGUCGUUCACGUUCACCUgc |
| BC042610 | EFNA4 | ephrin-A4 | 1945 | n371795 | AGAACAAAACUUCAGGGUUtt | AACCCUGAAGUUUUGUUCUtt |
| BC042610 | EFNA4 | ephrin-A4 | 1945 | n371794 | AGACAACCAACUCAGCUAAtt | UUAGCUGAGUUGGUUGUCUct |
| BC042610 | EFNA4 | ephrin-A4 | 1945 | n371793 | AGUCCGUCUUCAUAGAAGAtt | UCUUCUAUGAAGACGGACUct |
| NM_001962 | EFNA5 | ephrin-A5 | 1946 | s4508 | GGCCGAGAAUAUUUCUACAtt | UGUAGAAAUAUUCUCGGCCtg |
| NM_001962 | EFNA5 | ephrin-A5 | 1946 | s4507 | CGAUGUUAACGACAAAGUAtt | UACUUUGUCGUUAACAUCGaa |
| NM_001962 | EFNA5 | ephrin-A5 | 1946 | s4509 | GGUCCUGUCUAAAGCUCAAtt | UUGAGCUUUAGACAGGACCtt |
| NM_004429 | EFNB1 | ephrin-B1 | 1947 | s4511 | GGAAGGGCUUGGUGAUCUAtt | UAGAUCACCAAGCCCUUCCca |
| NM_004429 | EFNB1 | ephrin-B1 | 1947 | s4510 | UGACGGUCCUACUACUGAAtt | UUCAGUAGUAGGACCGUCAgg |
| NM_004429 | EFNB1 | ephrin-B1 | 1947 | s4512 | CUGUCUACAUCGUCCAAGAtt | UCUUGGACGAUGUAGACAGgg |
| NM_004093 | EFNB2 | ephrin-B2 | 1948 | s4515 | GGAAGAAGUUCGACAACAAtt | UUGUUGUCGAACUUCUUCCat |
| NM_004093 | EFNB2 | ephrin-B2 | 1948 | s4513 | CCAAAUCGAUAGUUUUAGAtt | UCUAAAACUAUCGAUUUGGaa |
| NM_004093 | EFNB2 | ephrin-B2 | 1948 | s4514 | GACAAGGACUGGUACUAUAtt | UAUAGUACCAGUCCUUGUCca |
| NM_004431 | EPHA2 | EPH receptor A2 | 1969 | s4564 | UGAUGAUCAUCACUGAGUAtt | UACUCAGUGAUGAUCAUCAtg |
| NM_004431 | EPHA2 | EPH receptor A2 | 1969 | s4566 | GUAUCUUCAUUGAGCUCAAtt | UUGAGCUCAAUGAAGAUACgc |
| NM_004431 | EPHA2 | EPH receptor A2 | 1969 | s4565 | GGAAGUACGAGGUCACUUAtt | UAAGUGACCUCGUACUUCCac |
| AK023333 | CTTN | cortactin | 2017 | n345349 | GCGCCCAGCUGAAACUUGAtt | UCAAGUUUCAGCUGGGCGCag |
| AK023333 | CTTN | cortactin | 2017 | n345348 | CCUGAGCAGUUGAUUUUCAtt | UGAAAAUCAACUGCUCAGGta |
| AK023333 | CTTN | cortactin | 2017 | n345347 | AAACCAGAGUUGUUAAAUUtt | AAUUUAACAACUCUGGUUUct |
| NM_005232 | EPHA1 | EPH receptor A1 | 2041 | s4721 | CCUCAGUAAUCACAAUUAUtt | AUAAUUGUGAUUACUGAGGta |
| NM_005232 | EPHA1 | EPH receptor A1 | 2041 | s4719 | GGUACCAGAUGGUUCUAGAtt | UCUAGAACCAUCUGGUACCgt |
| NM_005232 | EPHA1 | EPH receptor A1 | 2041 | s223490 | GAUGAAGAACGGUACCAGAtt | UCUGGUACCGUUCUUCAUCct |
| NM_182644 | EPHA3 | EPH receptor A3 | 2042 | s4724 | CGAGGUCAAAUACUAUGAAtt | UUCAUAGUAUUUGACCUCGta |
| NM_182644 | EPHA3 | EPH receptor A3 | 2042 | s4722 | GGAUGUACUGCAGUACAGAtt | UCUGUACUGCAGUACAUCCtt |
| NM_182644 | EPHA3 | EPH receptor A3 | 2042 | s4723 | CACCUGUCCUGACGAUUAAtt | UUAAUCGUCAGGACAGGUGat |
| NM_004440 | EPHA7 | EPH receptor A7 | 2045 | s4733 | CAGCUAAUGUUGGAUUGUUtt | AACAAUCCAACAUUAGCUGgt |
| NM_004440 | EPHA7 | EPH receptor A7 | 2045 | s4731 | GAUGGGCAAUUUACAGUCAtt | UGACUGUAAAUUGCCCAUCat |
| NM_004440 | EPHA7 | EPH receptor A7 | 2045 | s4732 | GGAACGGACCUACUCAACAtt | UGUUGAGUAGGUCCGUUCCct |
| NM_001006943 | EPHA8 | EPH receptor A8 | 2046 | s4736 | CCUCAAAAUCGACACCAUUtt | AAUGGUGUCGAUUUUGAGGaa |
| NM_001006943 | EPHA8 | EPH receptor A8 | 2046 | s4735 | GGACAUCAGUGACUCUGGAtt | UCCAGAGUCACUGAUGUCCca |
| NM_001006943 | EPHA8 | EPH receptor A8 | 2046 | s223498 | GUCUCAAGCUCAACACGGAtt | UCCGUGUUGAGCUUGAGACgc |
| NM_004442 | EPHB2 | EPH receptor B2 | 2048 | s4742 | AGAUGAUCCGCAAUCCCAAtt | UUGGGAUUGCGGAUCAUCUtg |
| NM_004442 | EPHB2 | EPH receptor B2 | 2048 | s4741 | ACAUCGAUCCUUUCACCUAtt | UAGGUGAAAGGAUCGAUGUag |
| NM_004442 | EPHB2 | EPH receptor B2 | 2048 | s4740 | GCGUGAUCCUGGACUAUGAtt | UCAUAGUCCAGGAUCACGCca |
| NM_004443 | EPHB3 | EPH receptor B3 | 2049 | s4743 | GGGUCUACGUGGAGCUCAAtt | UUGAGCUCCACGUAGACCCgc |
| NM_004443 | EPHB3 | EPH receptor B3 | 2049 | s4745 | CGAUCCUACCUACACCAGUtt | ACUGGUGUAGGUAGGAUCGga |
| NM_004443 | EPHB3 | EPH receptor B3 | 2049 | s4744 | GAAUGAAGGUUUAUAUUGAtt | UCAAUAUAAACCUUCAUUCca |
| NM_005246 | FER | fer (fps/fes related) tyrosine kinase | 2241 | s5110 | GAAUUAUGUCAGCAACGUAtt | UACGUUGCUGACAUAAUUCat |
| NM_005246 | FER | fer (fps/fes related) tyrosine kinase | 2241 | s5111 | GAACAACGGCUGCUAAAGAtt | UCUUUAGCAGCCGUUGUUCtg |
| NM_005246 | FER | fer (fps/fes related) tyrosine kinase | 2241 | s5109 | CACGAUCAGUUACAUCUAUtt | AUAGAUGUAACUGAUCGUGca |
| NM_002005 | FES | feline sarcoma oncogene | 2242 | s5114 | AGUGGGUGCUGAACCAUGAtt | UCAUGGUUCAGCACCCACUtg |
| NM_002005 | FES | feline sarcoma oncogene | 2242 | s5113 | CCUCAGCAAUCAGCAGACAtt | UGUCUGCUGAUUGCUGAGGtt |
| NM_002005 | FES | feline sarcoma oncogene | 2242 | s5112 | CCACGCUGGAGAUCCUUAAtt | UUAAGGAUCUCCAGCGUGGgt |
| NM_004475 | FLOT2 | flotillin 2 | 2319 | s5285 | ACAGUAAGGUCACAUCAGAtt | UCUGAUGUGACCUUACUGUtg |
| NM_004475 | FLOT2 | flotillin 2 | 2319 | s5286 | GACUAUAAACAGUACGUGUtt | ACACGUACUGUUUAUAGUCgg |
| NM_004475 | FLOT2 | flotillin 2 | 2319 | s5284 | CCAAGAUUGCUGACUCUAAtt | UUAGAGUCAGCAAUCUUGGtg |
| NM_000165 | GJA1 | gap junction protein, alpha 1, 43kDa | 2697 | s5757 | ACUAGCUGCUGGACAUGAAtt | UUCAUGUCCAGCAGCUAGUtt |
| NM_000165 | GJA1 | gap junction protein, alpha 1, 43kDa | 2697 | s5758 | GGCUAAUUACAGUGCAGAAtt | UUCUGCACUGUAAUUAGCCca |
| NM_000165 | GJA1 | gap junction protein, alpha 1, 43kDa | 2697 | s5759 | GAACCUACAUCAUCAGUAUtt | AUACUGAUGAUGUAGGUUCgc |
| NM_021954 | GJA3 | gap junction protein, alpha 3, 46kDa | 2700 | s5761 | GCAUGCUACUAGCAUCUUUtt | AAAGAUGCUAGUAGCAUGCat |
| NM_021954 | GJA3 | gap junction protein, alpha 3, 46kDa | 2700 | s5762 | CUGUUAGAAUGAUUGCUGAtt | UCAGCAAUCAUUCUAACAGta |
| NM_021954 | GJA3 | gap junction protein, alpha 3, 46kDa | 2700 | s5760 | CCACCCUACUAUGCGCACAtt | UGUGCGCAUAGUAGGGUGGga |
| NM_002060 | GJA4 | gap junction protein, alpha 4, 37kDa | 2701 | s5763 | ACUCGACCGUGGUGGGUAAtt | UUACCCACCACGGUCGAGUgc |
| NM_002060 | GJA4 | gap junction protein, alpha 4, 37kDa | 2701 | s5765 | GGCUUUGCCUGAGCACAGAtt | UCUGUGCUCAGGCAAAGCCac |
| NM_002060 | GJA4 | gap junction protein, alpha 4, 37kDa | 2701 | s5764 | CGACCGUGGUGGGUAAGAUtt | AUCUUACCCACCACGGUCGag |
| NM_005266 | GJA5 | gap junction protein, alpha 5, 40kDa | 2702 | s5767 | GCACUCUGCUCAACACCUAtt | UAGGUGUUGAGCAGAGUGCcc |
| NM_005266 | GJA5 | gap junction protein, alpha 5, 40kDa | 2702 | s5768 | CUAUCAUAGUGACAAGCGAtt | UCGCUUGUCACUAUGAUAGcc |
| NM_005266 | GJA5 | gap junction protein, alpha 5, 40kDa | 2702 | s5766 | CCGGUCAACUGUUACGUAUtt | AUACGUAACAGUUGACCGGgt |
| NM_004004 | GJB2 | gap junction protein, beta 2, 26kDa | 2706 | s5775 | AGAGUGAAUUUAAGGACAUtt | AUGUCCUUAAAUUCACUCUtt |
| NM_004004 | GJB2 | gap junction protein, beta 2, 26kDa | 2706 | s5777 | GUUAUUUGCUAAUUAGAUAtt | UAUCUAAUUAGCAAAUAACac |
| NM_004004 | GJB2 | gap junction protein, beta 2, 26kDa | 2706 | s5776 | GGACAUCGAGGAGAUCAAAtt | UUUGAUCUCCUCGAUGUCCtt |
| NM_001005752 | GJB3 | gap junction protein, beta 3, 31kDa | 2707 | s5779 | AGUUCACUAAGUUAUGCAAtt | UUGCAUAACUUAGUGAACUca |
| NM_001005752 | GJB3 | gap junction protein, beta 3, 31kDa | 2707 | s5778 | CAAGCGAAUUAACUAUCUAtt | UAGAUAGUUAAUUCGCUUGtt |
| NM_001005752 | GJB3 | gap junction protein, beta 3, 31kDa | 2707 | s5780 | CGACCUACCGAGAAGAAAAtt | UUUUCUUCUCGGUAGGUCGgg |
| NM_005268 | GJB5 | gap junction protein, beta 5, 31.1kDa | 2709 | s5783 | CGCAGAUCCAUGUCCCAAUtt | AUUGGGACAUGGAUCUGCGtg |
| NM_005268 | GJB5 | gap junction protein, beta 5, 31.1kDa | 2709 | s223727 | GGACUUCGACUGCAAUACUtt | AGUAUUGCAGUCGAAGUCCtt |

| | | | | | | |
|---|---|---|---|---|---|---|
| NM_005268 | GJB5 | gap junction protein, beta 5, 31.1kDa | 2709 | s5782 | GGACAUCGCCUUUCUCUAUtt | AUAGAGAAAGGCGAUGUCCac |
| NM_000201 | ICAM1 | intercellular adhesion molecule 1 | 3383 | s7087 | AGUCAACAGCUAAAACCUUtt | AAGGUUUUAGCUGUUGACUgc |
| NM_000201 | ICAM1 | intercellular adhesion molecule 1 | 3383 | s7088 | AGACUACAACAGGCCCAAAtt | UUUGGGCCUGUUGUAGUCUgt |
| NM_000201 | ICAM1 | intercellular adhesion molecule 1 | 3383 | s7086 | CGGAAGGUGUAUGAACUGAtt | UCAGUUCAUACACCUUCCGgt |
| NM_002230 | JUP | junction plakoglobin | 3728 | s7668 | GAUCAUGCGUAACUACAGUtt | ACUGUAGUUACGCAUGAUCtg |
| NM_002230 | JUP | junction plakoglobin | 3728 | s7666 | CUCUGUGCGUCUCAACUAUtt | AUAGUUGAGACGCACAGAGtt |
| NM_002230 | JUP | junction plakoglobin | 3728 | s7667 | CCAUCGGCUUGAUCAGGAAtt | UUCCUGAUCAAGCCGAUGGtt |
| NM_002444 | MSN | moesin | 4478 | s8985 | GCAGUACCAGGACACUAAAtt | UUUAGUGUCCUGGUACUGCag |
| NM_002444 | MSN | moesin | 4478 | s8984 | GGCUGAAACUCAAUAAGAAtt | UUCUUAUUGAGUUUCAGCCag |
| NM_002444 | MSN | moesin | 4478 | s8986 | GGUGUGAACUACUUCAGCAtt | UGCUGAAGUAGUUCACACCat |
| NM_005389 | PCMT1 | protein-L-isoaspartate (D-aspartate) O- | 5110 | s10130 | CCCACUAUGCAAAAUGUAAtt | UUACAUUUUGCAUAGUGGGag |
| NM_005389 | PCMT1 | protein-L-isoaspartate (D-aspartate) O- | 5110 | s10132 | CAAUCUCCGCAAAAAUGGAtt | UCCAUUUUUGCGGAGAUUGtg |
| NM_005389 | PCMT1 | protein-L-isoaspartate (D-aspartate) O- | 5110 | s10131 | GGAUUCUCCACAAUCAAUAtt | UAUUGAUUGUGGAGAAUCCat |
| NM_001042724 | PVRL2 | poliovirus receptor-related 2 (herpesvirus entry | 5819 | s11607 | CGCUGAGCAGGUCAUCUUUtt | AAAGAUGACCUGCUCAGCGcg |
| NM_001042724 | PVRL2 | poliovirus receptor-related 2 (herpesvirus entry | 5819 | s11606 | GCUAUGAUGACAACUGGUAtt | UACCAGUUGUCAUCAUAGCcg |
| NM_001042724 | PVRL2 | poliovirus receptor-related 2 (herpesvirus entry | 5819 | s11608 | GGCUAUGACUGGAGCACGAtt | UCGUGCUCCAGUCAUAGCCcg |
| NM_004162 | RAB5A | RAB5A, member RAS oncogene family | 5868 | s11678 | GGAAGAGGAGUAGACCUUAtt | UAAGGUCUACUCCUCUUCCtc |
| NM_004162 | RAB5A | RAB5A, member RAS oncogene family | 5868 | s11680 | GCAAGCAAGUCCUAACAUUtt | AAUGUUAGGACUUGCUUGCct |
| NM_004162 | RAB5A | RAB5A, member RAS oncogene family | 5868 | s11679 | CAAGCCUAGUGCUUCGUUUtt | AAACGAAGCACUAGGCUUGat |
| NM_006908 | RAC1 | ras-related C3 botulinum toxin substrate 1 (rho | 5879 | s11712 | ACAGAUUAAUUUUUCCAUAtt | UAUGGAAAAAUUAAUCUGUaa |
| NM_006908 | RAC1 | ras-related C3 botulinum toxin substrate 1 (rho | 5879 | s11711 | CUACUGUCUUUGACAAUUAtt | UAAUUGUCAAAGACAGUAGgg |
| NM_006908 | RAC1 | ras-related C3 botulinum toxin substrate 1 (rho | 5879 | s11713 | GGAACUAAACUUGAUCUUAtt | UAAGAUCAAGUUUAGUUCCca |
| NM_002906 | RDX | radixin | 5962 | s11899 | GAACUAUACAUGCGAAGAAtt | UUCUUCGCAUGUAUAGUUCat |
| NM_002906 | RDX | radixin | 5962 | s11900 | GGAAGAACGUGUAACCGAAtt | UUCGGUUACACGUUCUUCCtc |
| NM_002906 | RDX | radixin | 5962 | s11901 | GAUUCGACAAGGCAAUACAtt | UGUAUUGCCUUGUCGAAUCtg |
| NM_003025 | SH3GL1 | SH3-domain GRB2-like 1 | 6455 | s12796 | GCAAGGCGGUGACAGAAGUtt | ACUUCUGUCACCGCCUUGCtg |
| NM_003025 | SH3GL1 | SH3-domain GRB2-like 1 | 6455 | s12797 | GACUUUGACUACAAGAAGAtt | UCUUCUUGUAGUCAAAGUCca |
| NM_003025 | SH3GL1 | SH3-domain GRB2-like 1 | 6455 | s12798 | GCUUCAUCGUCUUUCCGAUtt | AUCGGAAAGACGAUGAAGCtg |
| NM_003257 | TJP1 | tight junction protein 1 (zona occludens 1) | 7082 | s14155 | GGAUGUUUAUCGUCGCAUUtt | AAUGCGACGAUAAACAUCCtt |
| NM_003257 | TJP1 | tight junction protein 1 (zona occludens 1) | 7082 | s14156 | CGAUCUCAUAAACUUCGUAtt | UACGAAGUUUAUGAGAUCGct |
| NM_003257 | TJP1 | tight junction protein 1 (zona occludens 1) | 7082 | s14157 | CUGUCAUACUUUGACCGAAtt | UUCGGUCAAAGUAUGACAGct |
| NM_006289 | TLN1 | talin 1 | 7094 | s14187 | GGAUUUUGGAGAUUACCAAtt | UUGGUAAUCUCCAAAAUCCag |
| NM_006289 | TLN1 | talin 1 | 7094 | s14186 | GCAGUUGACAGGACAUUCAtt | UGAAUGUCCUGUCAACUGCtg |
| NM_006289 | TLN1 | talin 1 | 7094 | s14185 | GGCUAUUACUCAGUACAGAtt | UCUGUACUGAGUAAUAGCCat |
| NM_001025234 | TSPAN4 | tetraspanin 4 | 7106 | s14216 | CGCCUACACGGACAAGAUUtt | AAUCUUGUCCGUGUAGGCGaa |
| NM_001025234 | TSPAN4 | tetraspanin 4 | 7106 | s14215 | CGGACAAGAUUGACAGGUAtt | UACCUGUCAAUCUUGUCCGtg |
| NM_001025234 | TSPAN4 | tetraspanin 4 | 7106 | s14217 | CCAACUACACUGACUGGUUtt | AACCAGUCAGUGUAGUUGGag |
| NM_006292 | TSG101 | tumor susceptibility gene 101 | 7251 | s14439 | GAAAAAGGGUCACCAGAAAtt | UUUCUGGUGACCCUUUUUCag |
| NM_006292 | TSG101 | tumor susceptibility gene 101 | 7251 | s14440 | CUGUCAAUGUUAUUACUCUtt | AGAGUAAUAACAUUGACAGtt |
| NM_006292 | TSG101 | tumor susceptibility gene 101 | 7251 | s14441 | GAGACCUAACUGUACGUGAtt | UCACGUACAGUUAGGUCUCtg |
| NM_001078 | VCAM1 | vascular cell adhesion molecule 1 | 7412 | s14759 | GGAGUUAAUUUGAUUGGGAtt | UCCCAAUCAAAUUAACUCCtt |
| NM_001078 | VCAM1 | vascular cell adhesion molecule 1 | 7412 | s14761 | GCAACUUGUGAAUCUAGGAtt | UCCUAGAUUCACAAGUUGCtg |
| NM_001078 | VCAM1 | vascular cell adhesion molecule 1 | 7412 | s14760 | GCAAGGUUCCUAGCGUGUAtt | UACACGCUAGGAACCUUGCag |
| NM_003373 | VCL | vinculin | 7414 | s14762 | GGUUGGUACUGCUAAUAAAtt | UUUAUUAGCAGUACCAACCgc |
| NM_003373 | VCL | vinculin | 7414 | s14763 | GCUUCAAUCAAAAUUCGAAtt | UUCGAAUUUUGAUUGAAGCag |
| NM_003373 | VCL | vinculin | 7414 | s14764 | GCUCGUAUCUUACUUAGGAtt | UCCUAAGUAAGAUACGAGCag |
| NM_003379 | EZR | ezrin | 7430 | s14796 | GGCUUUCCUUGGAGUGAAAtt | UUUCACUCCAAGGAAAGCCaa |
| NM_003379 | EZR | ezrin | 7430 | s14795 | CGUGGGAUGCUCAAAGAUAtt | UAUCUUUGAGCAUCCCACGgt |
| NM_003379 | EZR | ezrin | 7430 | s14797 | GGAAUCAACUAUUUCGAGAtt | UCUCGAAAUAGUUGAUUCCat |
| NM_004657 | SDPR | serum deprivation response | 8436 | s16008 | ACAUCGACUUGACUAUUGUtt | ACAAUAGUCAAGUCGAUGUtg |
| NM_004657 | SDPR | serum deprivation response | 8436 | s16006 | CGACUUGACUAUUGUGGAAtt | UUCCACAAUAGUCAAGUCGat |
| NM_004657 | SDPR | serum deprivation response | 8436 | s16007 | CAAAGAUCGUAUCUGUAGAtt | UCUACAGAUACGAUCUUUGtc |
| NM_003644 | GAS7 | growth arrest-specific 7 | 8522 | s16201 | AGACUAUGCGAAGAACUUAtt | UAAGUUCUUCGCAUAGUCUtc |
| NM_003644 | GAS7 | growth arrest-specific 7 | 8522 | s16202 | GAAUUCAUCCGGGAAAGGAtt | UCCUUUCCCGGAUGAAUUCtg |
| NM_003644 | GAS7 | growth arrest-specific 7 | 8522 | s16200 | GGCCCAGUCCAAAUGGUUUtt | AAACCAUUUGGACUGGGCCtg |
| NM_003941 | WASL | Wiskott-Aldrich syndrome-like | 8976 | s17134 | GCAGCUCUUUUAGAUCAAAtt | UUUGAUCUAAAAGAGCUGCtt |
| NM_003941 | WASL | Wiskott-Aldrich syndrome-like | 8976 | s17133 | GGAAUUGUGGGUGCAUUAAtt | UUAAUGCACCCACAAUUCCtg |
| NM_003941 | WASL | Wiskott-Aldrich syndrome-like | 8976 | s17132 | CGACAGGGUAUCCAACUAAtt | UUAGUUGGAUACCCUGUCGta |
| NM_024430 | PSTPIP2 | proline-serine-threonine phosphatase | 9050 | s17253 | CAAAAGAGCUUACAAUUCAtt | UGAAUUGUAAGCUCUUUUGtt |
| NM_024430 | PSTPIP2 | proline-serine-threonine phosphatase | 9050 | s17254 | CAGUUUGCUCUAUCAGUAAtt | UUACUGAUAGAGCAAACUGta |
| NM_024430 | PSTPIP2 | proline-serine-threonine phosphatase | 9050 | s17252 | CAUCGGCUAUGACAACAUUtt | AAUGUUGUCAUAGCCGAUGgt |
| NM_003978 | PSTPIP1 | proline-serine-threonine phosphatase | 9051 | s17257 | AGCUGUCGCUCUACAAGAAtt | UUCUUGUAGAGCGACAGCUtg |
| NM_003978 | PSTPIP1 | proline-serine-threonine phosphatase | 9051 | s17255 | GGAGAAGAGUCAGAACAAAtt | UUUGUUCUGACUCUUCUCCac |
| NM_003978 | PSTPIP1 | proline-serine-threonine phosphatase | 9051 | s17256 | CUACCAGAACUAUUACGAUtt | AUCGUAAUAGUUCUGGUAGgg |
| NM_012129 | CLDN12 | claudin 12 | 9069 | s17298 | CCAUCUGAACAAGAAGUUUtt | AAACUUCUUGUUCAGAUGGat |
| NM_012129 | CLDN12 | claudin 12 | 9069 | s17297 | GACACUACUUGGUACUCAUtt | AUGAGUACCAAGUAGUGUCgt |
| NM_012129 | CLDN12 | claudin 12 | 9069 | s17299 | CAUUCAACAGAAACGAGAAtt | UUCUCGUUUCUGUUGAAUGtg |
| NM_021195 | CLDN6 | claudin 6 | 9074 | s17309 | GGGAUGGGUUCGUACCUUUtt | AAAGGUACGAACCCAUCCCaa |
| NM_021195 | CLDN6 | claudin 6 | 9074 | s17311 | UGAGGAUAUUUAAAAUUCAtt | UGAAUUUUAAAUAUCCUCAgt |
| NM_021195 | CLDN6 | claudin 6 | 9074 | s17310 | GAUUGUCUUUGUCAUCUCAtt | UGAGAUGACAAAGACAAUCcc |
| NM_020384 | CLDN2 | claudin 2 | 9075 | s17314 | ACUCACCACUGGUGCCUGAtt | UCAGGCACCAGUGGUGAGUag |
| NM_020384 | CLDN2 | claudin 2 | 9075 | s17312 | GGAUCCUACGGGACUUCUAtt | UAGAAGUCCCGUAGGAUCCca |
| NM_020384 | CLDN2 | claudin 2 | 9075 | s225076 | GCUCUUUACUUGGGCAUUAtt | UAAUGCCCAAGUAAAGAGCct |
| NM_021101 | CLDN1 | claudin 1 | 9076 | s17316 | GACUCCUUGCUGAAUCUGAtt | UCAGAUUCAGCAAGGAGUCaa |
| NM_021101 | CLDN1 | claudin 1 | 9076 | s17317 | ACCUCUUACCCAACACCAAtt | UUGGUGUUGGGUAAGAGGUtg |
| NM_021101 | CLDN1 | claudin 1 | 9076 | s17315 | CAAUAGAAUCGUUCAAGAAtt | UUCUUGAACGAUUCUAUUGcc |
| NM_020982 | CLDN9 | claudin 9 | 9080 | s194981 | ACCCUGGAGUUGUCAGGGAtt | UCCCUGACAACUCCAGGGUtg |
| NM_020982 | CLDN9 | claudin 9 | 9080 | s194980 | GCAGCUCGUUGGCUCCAAAtt | UUUGGAGCCAACGAGCUGCtg |
| NM_020982 | CLDN9 | claudin 9 | 9080 | s194979 | CCCUCGAGUUGGAAAUAAAtt | UUUAUUUCCAACUCGAGGGtg |
| NM_004712 | HGS | hepatocyte growth factor-regulated tyrosine | 9146 | s17480 | CGUCUUUCCAGAAUUCAAAtt | UUUGAAUUCUGGAAAGACGtg |
| NM_004712 | HGS | hepatocyte growth factor-regulated tyrosine | 9146 | s17481 | UGGAAUCUGUGGUAAAGAAtt | UUCUUUACCACAGAUUCCAtg |

| | | | | | | |
|---|---|---|---|---|---|---|
| NM_004712 | HGS | hepatocyte growth factor-regulated tyrosine | 9146 | s17482 | CACGGUAUCUCAACCGGAAtt | UUCCGGUUGAGAUACCGUGcg |
| NM_004240 | TRIP10 | thyroid hormone receptor interactor 10 | 9322 | s17816 | GAACCUCAGUGUCCGUGUAtt | UACACGGACACUGAGGUUCtc |
| NM_004240 | TRIP10 | thyroid hormone receptor interactor 10 | 9322 | s17815 | GGAAGAACGCAGUCGUGAAtt | UUCACGACUGCGUUCUUCCaa |
| NM_004240 | TRIP10 | thyroid hormone receptor interactor 10 | 9322 | s17814 | GGAGAAUAGUAAGCGUAAAtt | UUUACGCUUACUAUUCUCCag |
| NM_004869 | VPS4B | vacuolar protein sorting 4 homolog B (S. | 9525 | s18274 | CGACCAAAUGUGAAAUGGAtt | UCCAUUUCACAUUUGGUCGtt |
| NM_004869 | VPS4B | vacuolar protein sorting 4 homolog B (S. | 9525 | s18273 | GCGAAGAUUUGAGAAACGAtt | UCGUUUCUCAAAUCUUCGCct |
| NM_004869 | VPS4B | vacuolar protein sorting 4 homolog B (S. | 9525 | s18272 | CACGUAGAAUUAAGACGGAtt | UCCGUCUUAAUUCUACGUGcg |
| NM_014751 | MTSS1 | metastasis suppressor 1 | 9788 | s18916 | CCAGCUUCGGACAACAGUAtt | UACUGUUGUCCGAAGCUGGga |
| NM_014751 | MTSS1 | metastasis suppressor 1 | 9788 | s18917 | CCAUCAUCAGCGACAUGAAtt | UUCAUGUCGCUGAUGAUGGtc |
| NM_014751 | MTSS1 | metastasis suppressor 1 | 9788 | s18915 | CCAUCUCGGAAGAUCUAAAtt | UUUAGAUCUUCCGAGAUGGtc |
| NM_001080383 | GJC1 | gap junction protein, gamma 1, 45kDa | 10052 | s19541 | GGAGGAGAAUCCAUCUAUUtt | AAUAGAUGGAUUCUCCUCCta |
| NM_001080383 | GJC1 | gap junction protein, gamma 1, 45kDa | 10052 | s19543 | UGUCUGUUAUGAUGCGUUUtt | AAACGCAUCAUAACAGACAtt |
| NM_001080383 | GJC1 | gap junction protein, gamma 1, 45kDa | 10052 | s223072 | GAUUCACAACCAUUCCACAtt | UGUGGAAUGGUUGUGAAUCtc |
| NM_005803 | FLOT1 | flotillin 1 | 10211 | s19915 | AGAGAGAUUACGAACUGAAtt | UUCAGUUCGUAAUCUCUCUgt |
| NM_005803 | FLOT1 | flotillin 1 | 10211 | s19914 | GCAUCAGUGUGGUUAGCUAtt | UAGCUAACCACACUGAUGCcc |
| NM_005803 | FLOT1 | flotillin 1 | 10211 | s19913 | GCAGAGAAGUCCCAACUAAtt | UUAGUUGGGACUUCUCUGCct |
| NM_017451 | BAIAP2 | BAI1-associated protein 2 | 10458 | s20464 | AGAAGUACUCGGACAAGGAtt | UCCUUGUCCGAGUACUUCUga |
| NM_017451 | BAIAP2 | BAI1-associated protein 2 | 10458 | s20465 | CAGAAUCAGCUGGAAGAAAtt | UUUCUUCCAGCUGAUUCUGga |
| NM_017451 | BAIAP2 | BAI1-associated protein 2 | 10458 | s20463 | AGACCGCACUGACAGAGGAtt | UCCUCUGUCAGUGCGGUCUtg |
| NM_006783 | GJB6 | gap junction protein, beta 6, 30kDa | 10804 | s21231 | AGACAUAGAGGACAUUAAAtt | UUUAAUGUCCUCUAUGUCUtt |
| NM_006783 | GJB6 | gap junction protein, beta 6, 30kDa | 10804 | s21232 | GAGAGUAAGCAGAAUGAAAtt | UUUCAUUCUGCUUACUCUCct |
| NM_006783 | GJB6 | gap junction protein, beta 6, 30kDa | 10804 | s21233 | GCAGCAUCUUUUUCCGAAUtt | AUUCGGAAAAAGAUGCUGCtg |
| NM_007229 | PACSIN2 | protein kinase C and casein kinase substrate in | 11252 | s229822 | ACCCAUCCCUCAACCCUGAtt | UCAGGGUUGAGGGAUGGGUct |
| NM_007229 | PACSIN2 | protein kinase C and casein kinase substrate in | 11252 | s22216 | AGAUGUUCUUAAGACCAAAtt | UUUGGUCUUAAGAACAUCUtg |
| NM_007229 | PACSIN2 | protein kinase C and casein kinase substrate in | 11252 | s22215 | GGAGGUUCAGAAGCACCUAtt | UAGGUGCUUCUGAACCUCCag |
| NM_007241 | SNF8 | SNF8, ESCRT-II complex subunit, homolog (S. | 11267 | s22248 | GCCCAGAUGUCAAAGCAGUtt | ACUGCUUUGACAUCUGGGCta |
| NM_007241 | SNF8 | SNF8, ESCRT-II complex subunit, homolog (S. | 11267 | s22247 | CAACAGGUGUUGAAGGGAAtt | UUCCCUUCAACACCUGUUGat |
| NM_007241 | SNF8 | SNF8, ESCRT-II complex subunit, homolog (S. | 11267 | s22249 | CGAACUAGGUGUCCAAAUUtt | AAUUUGGACACCUAGUUCGta |
| NM_015033 | FNBP1 | formin binding protein 1 | 23048 | s22914 | CCGAAUCAAUUGAUCAGAAtt | UUCUGAUCAAUUGAUUCGGct |
| NM_015033 | FNBP1 | formin binding protein 1 | 23048 | s22915 | GGAUUUUGACGACGAGUUUtt | AAACUCGUCGUCAAAAUCCgt |
| NM_015033 | FNBP1 | formin binding protein 1 | 23048 | s22916 | CAACCUAAAAAGAACUCGAtt | UCGAGUUCUUUUUAGGUUGgt |
| NM_015221 | DNMBP | dynamin binding protein | 23268 | s23437 | GACCAUUCCCAGUCUAAAAtt | UUUUAGACUGGGAAUGGUCac |
| NM_015221 | DNMBP | dynamin binding protein | 23268 | s23438 | CCGAGUAAUUCAGAGAGUAtt | UACUCUCUGAAUUACUCGGat |
| NM_015221 | DNMBP | dynamin binding protein | 23268 | s23439 | CGUUACCCGCUGUUGCUAAtt | UUAGCAACAGCGGGUAACGca |
| NM_001042758 | SRGAP2 | SLIT-ROBO Rho GTPase activating protein 2 | 23380 | s23693 | AGUACAUCGUGGUCCAAGAtt | UCUUGGACCACGAUGUACUga |
| NM_001042758 | SRGAP2 | SLIT-ROBO Rho GTPase activating protein 2 | 23380 | s23692 | GCAUGGAGGAUUACUGUGAtt | UCACAGUAAUCCUCCAUGCtt |
| NM_001042758 | SRGAP2 | SLIT-ROBO Rho GTPase activating protein 2 | 23380 | s23691 | CUCUGAUUAUCAUGAGAUAtt | UAUCUCAUGAUAAUCAGAGtg |
| NM_012120 | CD2AP | CD2-associated protein | 23607 | s24191 | CGAACUUAAUGGUAAAGAAtt | UUCUUUACCAUUAAGUUCGcc |
| NM_012120 | CD2AP | CD2-associated protein | 23607 | s24192 | CUAUGAAGGUACUAAUGAAtt | UUCAUUAGUACCUUCAUAGgc |
| NM_012120 | CD2AP | CD2-associated protein | 23607 | s24193 | CUGGAACAGUGUACCCAAAtt | UUUGGGUACACUGUUCCAGaa |
| NM_015480 | PVRL3 | poliovirus receptor-related 3 | 25945 | s24804 | GGAAAUUGGUUUGUAGGAAtt | UUCCUACAAACCAAUUUCCat |
| NM_015480 | PVRL3 | poliovirus receptor-related 3 | 25945 | s24803 | CCAUUGACUUUCAAUUAUUtt | AAUAAUUGAAAGUCAAUGGat |
| NM_015480 | PVRL3 | poliovirus receptor-related 3 | 25945 | s24802 | GGAAUAUUCUGCUAUAGGAtt | UCCUAUAGCAGAAUAUUCCag |
| NM_015569 | DNM3 | dynamin 3 | 26052 | s25018 | GACUCCUACAUGUCCAUUAtt | UAAUGGACAUGUAGGAGUCta |
| NM_015569 | DNM3 | dynamin 3 | 26052 | s25017 | GCAUUUGAAGCGAUAGUCAtt | UGACUAUCGCUUCAAAUGCca |
| NM_015569 | DNM3 | dynamin 3 | 26052 | s25019 | GUGUGGAUCUGGUAAUACAtt | UGUAUUACCAGAUCCACACtc |
| NM_016223 | PACSIN3 | protein kinase C and casein kinase substrate in | 29763 | s26562 | GGAUGAGGAGUGGUCAGAUtt | AUCUGACCACUCCUCAUCCtg |
| NM_016223 | PACSIN3 | protein kinase C and casein kinase substrate in | 29763 | s26563 | CAAGAAAAGCUACCACGCAtt | UGCGUGGUAGCUUUUCUUGga |
| NM_016223 | PACSIN3 | protein kinase C and casein kinase substrate in | 29763 | s26564 | CAAUCAGCCGGAAAGAGAAtt | UUCUCUUUCCGGCUGAUUGtc |
| NM_018482 | ASAP1 | ArfGAP with SH3 domain, ankyrin repeat and | 50807 | s27099 | GGAUAUCAGUAUUGACAAAtt | UUUGUCAAUACUGAUAUCCat |
| NM_018482 | ASAP1 | ArfGAP with SH3 domain, ankyrin repeat and | 50807 | s27101 | CACAAGUUCUUGAUAAGUUtt | AACUUAUCAAGAACUUGUGca |
| NM_018482 | ASAP1 | ArfGAP with SH3 domain, ankyrin repeat and | 50807 | s27100 | GCACCGCGUUUGUCAAGUUtt | AACUUGACAAACGCGGUGCca |
| NM_016946 | F11R | F11 receptor | 50848 | s27150 | GCCUAGUGCCCGAAGUGAAtt | UUCACUUCGGGCACUAGGCtg |
| NM_016946 | F11R | F11 receptor | 50848 | s27151 | CCAUCCAAGCCUACAGUUAtt | UAACUGUAGGCUUGGAUGGag |
| NM_016946 | F11R | F11 receptor | 50848 | s27152 | GGAUAGUGAUGCCUACGAAtt | UUCGUAGGCAUCACUAUCCca |
| NM_016208 | VPS28 | vacuolar protein sorting 28 homolog (S. | 51160 | s27577 | AAUCAGCUCUAUUGACGAAtt | UUCGUCAAUAGAGCUGAUUtc |
| NM_016208 | VPS28 | vacuolar protein sorting 28 homolog (S. | 51160 | s27578 | GGAACAAGCCGGAGCUGUAtt | UACAGCUCCGGCUUGUUCCca |
| NM_016208 | VPS28 | vacuolar protein sorting 28 homolog (S. | 51160 | s27579 | GAAGUGAAGUUGUACAAGAtt | UCUUGUACAACUUCACUUCct |
| NM_016580 | PCDH12 | protocadherin 12 | 51294 | s27870 | CAUACAUGUUGCUAACCAAtt | UUGGUUAGCAACAUGUAUGtg |
| NM_016580 | PCDH12 | protocadherin 12 | 51294 | s27869 | CCAACUACCGUGACAAUGUtt | ACAUUGUCACGGUAGUUGGtg |
| NM_016580 | PCDH12 | protocadherin 12 | 51294 | s27871 | CUGGUACAGUGAUCGGGAAtt | UUCCCGAUCACUGUACCAGat |
| NM_001005753 | VPS24 | vacuolar protein sorting 24 homolog (S. | 51652 | s28475 | GCUGGGAUCAUAGAGGAGAtt | UCUCCUCUAUGAUCCCAGCct |
| NM_001005753 | VPS24 | vacuolar protein sorting 24 homolog (S. | 51652 | s28474 | UGAGGGAGUUGUCCAAAGAtt | UCUUUGGACAACUCCCUCAtg |
| NM_001005753 | VPS24 | vacuolar protein sorting 24 homolog (S. | 51652 | s28473 | GCAUGGACGAUCAGGAAGAtt | UCUUCCUGAUCGUCCAUGCtt |
| NM_017737 | FNBP1L | formin binding protein 1-like | 54874 | s29645 | GGCACAACAGAGUUAUGAAtt | UUCAUAACUCUGUUGUGCCtt |
| NM_017737 | FNBP1L | formin binding protein 1-like | 54874 | s29646 | GGACUAUCAGUGCAUCCAAtt | UUGGAUGCACUGAUAGUCCca |
| NM_017737 | FNBP1L | formin binding protein 1-like | 54874 | s29644 | GGACGAACGAAGGACUAUUtt | AAUAGUCCUUCGUUCGUCCat |
| NM_020145 | SH3GLB2 | SH3-domain GRB2-like endophilin B2 | 56904 | s32358 | AGACAGAGUUUGACCGGCAtt | UGCCGGUCAAACUCUGUCUgg |
| NM_020145 | SH3GLB2 | SH3-domain GRB2-like endophilin B2 | 56904 | s32357 | CGACGGUGCCUGACUUUCAtt | UGAAAGUCAGGCACCGUCGtg |
| NM_020145 | SH3GLB2 | SH3-domain GRB2-like endophilin B2 | 56904 | s32359 | GGAAGACACUGAUCAAGGUtt | ACCUUGAUCAGUGUCUUCCca |
| NM_020762 | SRGAP1 | SLIT-ROBO Rho GTPase activating protein 1 | 57522 | s33218 | GCACUAAGGAUCAUCAACAtt | UGUUGAUGAUCCUUAGUGCtt |
| NM_020762 | SRGAP1 | SLIT-ROBO Rho GTPase activating protein 1 | 57522 | s33219 | GGUCGGUCCUUAUAGUGAUtt | AUCACUAUAAGGACCGACCtg |
| NM_020762 | SRGAP1 | SLIT-ROBO Rho GTPase activating protein 1 | 57522 | s33217 | GGAUAUGGAUGAUACGUUUtt | AAACGUAUCAUCCAUAUCCtg |
| NM_021219 | JAM2 | junctional adhesion molecule 2 | 58494 | s33856 | GCUACGAUGUCAAGACAAAtt | UUUGUCUUGACAUCGUAGCtc |
| NM_021219 | JAM2 | junctional adhesion molecule 2 | 58494 | s33857 | GCAUCCGUUUGCUAGAAAAtt | UUUUCUAGCAAACGGAUGCca |
| NM_021219 | JAM2 | junctional adhesion molecule 2 | 58494 | s33855 | CAGUCACUCUGGAAGUAUUtt | AAUACUUCCAGAGUGACUGta |
| NM_015059 | TLN2 | talin 2 | 83660 | s38090 | GCUGCUUGUGGAUUCGCUAtt | UAGCGAAUCCACAAGCAGCtt |
| NM_015059 | TLN2 | talin 2 | 83660 | s38088 | GGAGUAGAUGAAAACGAAAtt | UUUCGUUUUCAUCUACUCCtt |
| NM_015059 | TLN2 | talin 2 | 83660 | s38089 | CGACGAAUCCAAACACGAAtt | UUCGUGUUUGGAUUCGUCGac |
| NM_032801 | JAM3 | junctional adhesion molecule 3 | 83700 | s38127 | AGUUAACUGUGCAAGUGAAtt | UUCACUUGCACAGUUAACUcg |
| NM_032801 | JAM3 | junctional adhesion molecule 3 | 83700 | s38129 | GGAGUUAACUACAUCCGCAtt | UGCGGAUGUAGUUAACUCCat |

| | | | | | | |
|---|---|---|---|---|---|---|
| NM_032801 | JAM3 | junctional adhesion molecule 3 | 83700 | s38128 | GCUACUUCAUCAACAAUAAtt | UUAUUGUUGAUGAAGUAGCca |
| NM_138383 | MTSS1L | metastasis suppressor 1-like | 92154 | s40868 | CGGCAUGAGAUCAAAAAGAtt | UCUUUUUGAUCUCAUGCCGgg |
| NM_138383 | MTSS1L | metastasis suppressor 1-like | 92154 | s40867 | GCCCGAGUUUGACAAGUCAtt | UGACUUGUCAAACUCGGGCgg |
| NM_138383 | MTSS1L | metastasis suppressor 1-like | 92154 | s40869 | CAAACAUACUCACCCAGUUtt | AACUGGGUGAGUAUGUUUGgg |
| NM_001039724 | NOSTRIN | nitric oxide synthase trafficker | 115677 | s41845 | GGAGCAUACUCAUAGCUAUtt | AUAGCUAUGAGUAUGCUCCtt |
| NM_001039724 | NOSTRIN | nitric oxide synthase trafficker | 115677 | s41847 | GGCGGGUUAUUCUACCAGAtt | UCUGGUAGAAUAACCCGCCat |
| NM_001039724 | NOSTRIN | nitric oxide synthase trafficker | 115677 | s41846 | CUGAGUUCCUGUUAACGGAtt | UCCGUUAACAGGAACUCAGat |
| NM_152219 | GJD3 | gap junction protein, delta 3, 31.9kDa | 125111 | s42859 | GCAUCAUCCGAUGGAAAUAtt | UAUUUCCAUCGGAUGAUGCtg |
| NM_152219 | GJD3 | gap junction protein, delta 3, 31.9kDa | 125111 | s42857 | CUGCAACCGUGCACACGAAtt | UUCGUGUGCACGGUUGCAGcg |
| NM_152219 | GJD3 | gap junction protein, delta 3, 31.9kDa | 125111 | s42858 | CGGUGCUGUUCGUCGUCUAtt | UAGACGACGAACAGCACCGgg |
| NM_153212 | GJB4 | gap junction protein, beta 4, 30.3kDa | 127534 | s43186 | CAGUAUAUGUCAAACCUCUtt | AGAGGUUUGACAUAUACUGtt |
| NM_153212 | GJB4 | gap junction protein, beta 4, 30.3kDa | 127534 | s43185 | ACGACAACCUGAGCAAGAAtt | UUCUUGCUCAGGUUGUCGUac |
| NM_153212 | GJB4 | gap junction protein, beta 4, 30.3kDa | 127534 | s43184 | GCCUCUACAAGGAUUAUGAtt | UCAUAAUCCUUGUAGAGGCgg |
| NM_177986 | DSG4 | desmoglein 4 | 147409 | s44997 | GCUCCAGUCUUUUCGCAAAtt | UUUGCGAAAAGACUGGAGCgt |
| NM_177986 | DSG4 | desmoglein 4 | 147409 | s44999 | CGGCAAUCCUUACGGCUAAtt | UUAGCCGUAAGGAUUGCCGag |
| NM_177986 | DSG4 | desmoglein 4 | 147409 | s44998 | GUUCGGAACUGAUACGAUUtt | AAUCGUAUCAGUUCCGAACtt |
| NM_012232 | PTRF | polymerase I and transcript release factor | 284119 | s49506 | GUCGGGAUCUCAGAGGAAAtt | UUUCCUCUGAGAUCCCGACtt |
| NM_012232 | PTRF | polymerase I and transcript release factor | 284119 | s49507 | CAUCUCUACUAAGCGAAAAtt | UUUUCGCUUAGUAGAGAUGgg |
| NM_012232 | PTRF | polymerase I and transcript release factor | 284119 | s49508 | CGAGCAAUACGGUGAGCAAtt | UUGCUCACCGUAUUGCUCGtg |
| | OCLN | | | s446558 | AGACUAUGAUAGACAGAAAtt | UUUCUGUCUAUCAUAGUCUcc |
| | OCLN | | | s446559 | AGAACUUGAUGAGAUCAAUtt | AUUGAUCUCAUCAAGUUCUga |
| | OCLN | | | s446560 | AGAAUUGGAUGACUAUAGAtt | UCUAUAGUCAUCCAAUUCUtt |
